# Loading of extracellular vesicles with nucleic acids via hybridization with sponge-like lipid nanoparticles

**DOI:** 10.1101/2024.04.10.588678

**Authors:** Johannes Bader, Pascal Rüedi, Valeria Mantella, Silvana Geisshüsler, Finn Brigger, Bilal M. Qureshi, Jaime Ortega Arroyo, Elita Montanari, Jean-Christophe Leroux

## Abstract

The translation of cell-derived extracellular vesicles (EVs) into biogenic gene delivery systems is limited by relatively inefficient loading strategies. In this work, we describe the loading of various nucleic acids into small EVs *via* their spontaneous hybridization with preloaded non-lamellar liquid crystalline lipid nanoparticles (LCNPs) under physiological conditions, forming hybrid EVs (HEVs). We correlate LCNPs’ topological characteristics with their propensity to fuse/aggregate with EVs and found that sponge (L_3_) phases at pH 7.4 were particularly suitable to induce a controlled hybridization process. State-of-the-art single-particle analysis techniques revealed that L_3_-based LCNPs interact with various EV subpopulations and that around 40% of HEVs were loaded with the genetic cargo. Importantly, this study demonstrates that EV membrane proteins remain accessible on HEV surfaces, with their intrinsic enzymatic activity unaffected after the hybridization process. Finally, HEVs showed *in vitro* improved transfection efficiencies compared to unhybridized LCNPs. In summary, this versatile platform holds potential for loading various nucleic acid molecules into native EVs and may help developing EV-based therapeutics.

**Teaser:** Topology of lipid nanoparticles influences their hybridization behavior with extracellular vesicles and produces novel biogenic gene delivery systems.

## Introduction

In recent years, RNA therapeutics, such as mRNA-based vaccines and siRNA drugs, have received increasing interest, resulting in the approval of groundbreaking medicines like the COVID-19 vaccines, patisiran/Onpattro^®^ or nusinersen/Spinraza^®^ (*1*). While oligonucleotides can be chemically modified to enhance their “drug-likeness”, and are often administered naked or conjugated with targeting motifs (*e.g.*, N-acetylgalactosamine), larger nucleic acids (NAs) such as mRNA typically require encapsulation inside a nanocarrier to facilitate their intracellular delivery (*2–4*). Lipid nanoparticles (LNPs) have emerged as clinically relevant drug delivery systems for NAs owing to their ease of manufacturing, high NA loading efficiencies and generally good safety profile (*5*). However, the LNP field currently faces two obstacles that are slowing down its advancement. One limitation is that systemically injected LNPs lack tissue-specific targeting, accumulating mainly in organs of the mononuclear phagocyte system (*e.g.*, liver), necessitating the administration of high doses when other tissues are treated (*2, 6*). Second, LNPs have been linked to unwanted immune reactions and accelerated blood clearance after repeated administrations, underlining the need for the development of safer and more efficacious LNP formulations (7–9).

Extracellular vesicles (EVs), which are naturally occurring cell-produced vesicles, may offer a promising avenue to address some of the limitations of LNPs (*10*). They transport biomolecules such as proteins and NAs through the extracellular space, serving as important mediators of cell-cell communication (*11, 12*). Clinical trials with EVs (*e.g.*, from mesenchymal stem cells (MSCs)) have highlighted their good tolerability (*10, 13–15*), and there is evidence that EVs can deliver NAs more safely and effectively compared to LNPs (*16–19*). Nevertheless, achieving efficient loading of therapeutic NA doses into the lumen of small EVs (< 200 nm) remains challenging.

Genetic manipulation of EV producer cell lines is a widely used method to introduce protein or NA-based cargo into EVs (*20*). However, this approach has limitations when working with native EVs obtained from primary cells or patient-derived biofluids, and is restricted to cell-expressible NAs, excluding chemically modified ones (*10, 21*). Post-isolation loading methods are, therefore, more versatile. They usually temporarily disrupt EV membranes through physical or chemical stimuli, and include electroporation (*22–24*), sonication (*25*), extrusion (*26*), transmembrane pH gradients (*27*), and heat shocks (*28*). Other strategies encompass hydrophobic modification of NAs (*e.g.*, with cholesterol) (*29, 30*) for surface attachment onto EVs or interaction/hybridization with other nanocarriers (*e.g.*, complexes or liposomes) containing lipid-, polymer- or protein-based materials (*e.g.*, Lipofectamine^TM^, Exo-Fect^TM^ and poly(ethylenimine) (PEI)) (*31–36*). Despite over a decade of research on exogenous loading of EVs, a lack of standardized reporting of protocols and insufficient characterization of the engineered EV-based products have hindered their clinical translation (*37*). In fact, it has now been shown that several exogenous loading techniques and reagents can negatively affect the physical and functional properties of EVs (*24, 38–40*).

Motivated by the existing limitations of both fields, we report an exogenous loading strategy for EVs that accomplishes two primary goals: (1) avoids the use of harsh mechanical or chemical treatments; thereby preserving at least partially the functional and biological properties of EVs; (2) efficiently encapsulates NAs within EVs. In this context, we propose the use of non-lamellar liquid crystalline lipid nanoparticles (LCNPs) to transfect native MSC-derived EVs with various types of NA cargos, such as siRNA, mRNA and RNA aptamers. Using single-particle characterization techniques, we study the correlation between the physicochemical properties/topology of lipid nanocarriers and their propensity to fuse/aggregate with small EVs. With a state-of-the-art optofluidic platform we demonstrate that MSC-EV membrane proteins are located on surfaces of the formed hybrid EV (HEV) particles and their intrinsic enzymatic activity is not altered during the process. Finally, we show that the hybridization with EVs can increase the knockdown/expression efficiencies of siRNA- and mRNA-loaded LCNPs, particularly under challenging cell culture conditions.

## Results

### Production and characterization of LCNPs

Inspired by early studies in the 1990’s and 2000’s exploring non-lamellar phases and their role in promoting fusion stalk and endosomal escape in cationic liposomes, we engineered a type of non-lamellar LNP (*41–45*). We hypothesized that non-lamellar lyotropic liquid crystalline (LLC) domains in LNPs may in a similar way induce fusion processes with EVs. To create the LCNPs, the ionizable lipid heptatriaconta-6,9,28,31-tetraen-19-yl-4-(dimethylamino)butanoate (DLin-MC3-DMA (MC3)) was combined with the helper lipid 1,2-dioleoyl-*sn*-glycero-3-phosphoethanolamine (DOPE), a lipid known to induce inverted hexagonal (H_II_) phases by lowering the spontaneous membrane curvature (*C_0_*) (*45*). To obtain stable nanostructures in an aqueous environment, the non-ionic surfactant poly(ethylene glycol) 12-hydroxystearate (Kolliphor^®^ HS 15 (HS15)) was added to the mixture. HS15 was selected instead of traditional PEG-lipids, as the latter could potentially interfere with the fusion process between LCNPs and EVs (*46*). The LCNPs were produced *via* microfluidics by mixing the lipid/surfactant combination in ethanol with the NA-containing aqueous phase. An siRNA targeting the green fluorescent protein (GFP) mRNA (siGFP) was first loaded into LCNPs. Small-angle X-ray scattering (SAXS) and cryogenic transmission electron microscopy (cryo-TEM) were employed to probe the structural properties of LCNPs and assess the effect of replacing the inverse cone-shaped lipid DOPE with the cylindrical lipid 1,2-dioleoyl-*sn*-glycero-3-phosphocholine (DOPC) or the inverse cone-shaped MC3 with the cylindrical fixed cationic lipid 1,2-di-*O*-octadecenyl-3-trimethylammonium propane (DOTMA) (Fig. 1).

**Fig. 1.**
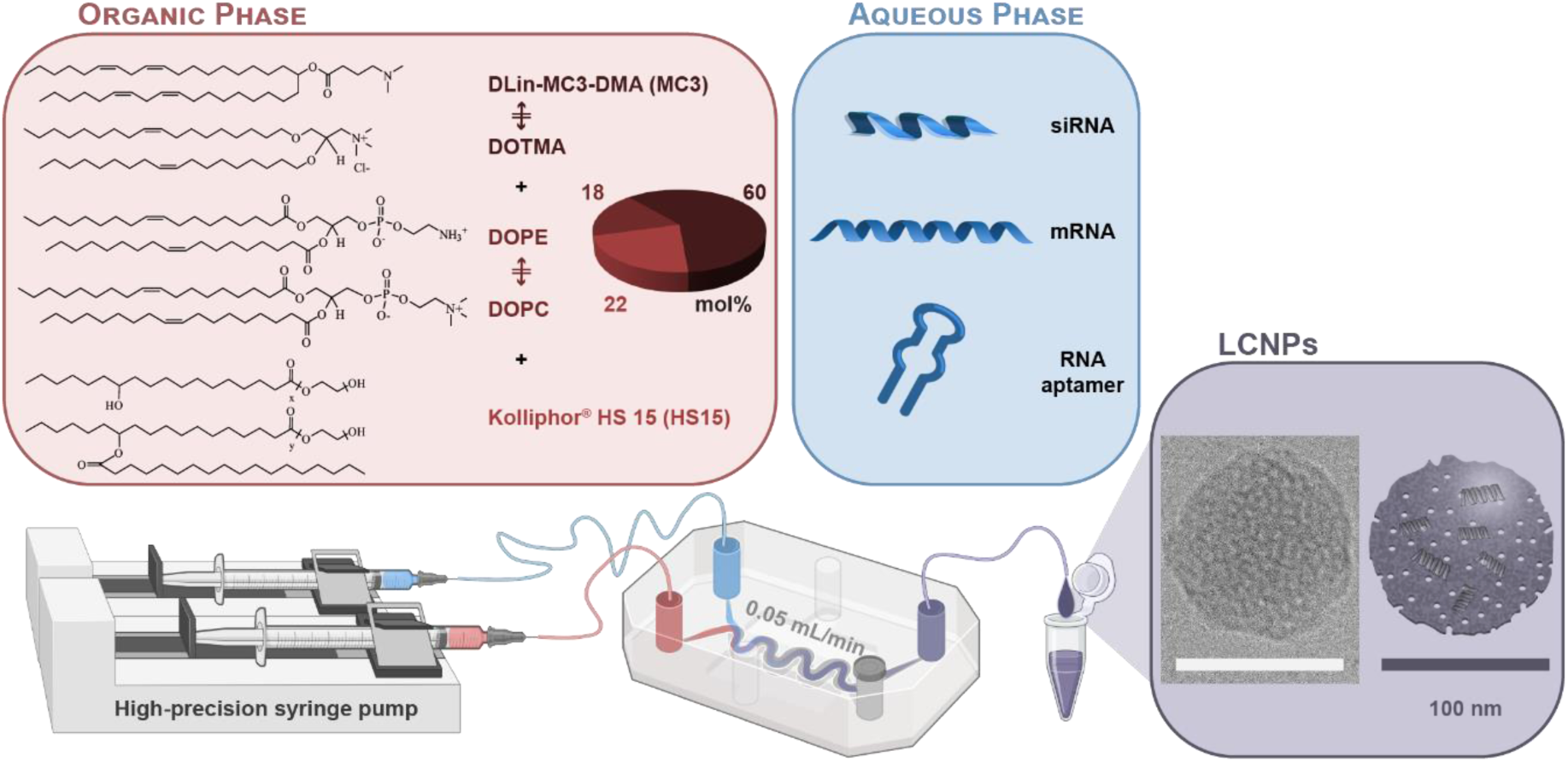
Formulation composition and microfluidic production process of LCNPs. The organic phase is composed of a tertiary mixture including the ionizable lipid MC3 (60 mol%), along with nearly equimolar proportions of the helper lipid DOPE (18 mol%) and the non-ionic surfactant HS15 (22 mol%) in ethanol. The cylindrical lipids DOPC and DOTMA were employed as substitutes for the inverted cone-shaped lipids DOPE and MC3, respectively. The aqueous phase contains the NAs (*i.e.*, siRNA, mRNA and RNA aptamer) dissolved in an acetate buffer (125 mM, pH 4). LCNPs are produced *via* microfluidic mixing of the lipid/surfactant-containing organic phase with the NA-containing aqueous phase utilizing a staggered diffusion mixer chip and a high-precision syringe pump (0.05 mL/min). The cryo-TEM image shows a representative siRNA-loaded LCNP (N/P 25); scale bar 100 nm. Created with BioRender.com.

LCNPs loaded with the siRNA at an N/P ratio of 25 had a hydrodynamic diameter (d_h_) of 115 nm at pH 7.4 as determined by dynamic light scattering (DLS) (Fig. 2A). Without the siRNA, LCNPs exhibited a d_h_ of 100 nm. In both cases, the d_h_ increased by approximately 40 nm under acidic conditions (pH 5). Similarly, zeta (ζ) potentials increased at pH 5 (Fig. 2A). At pH 7.4, siRNA-loaded LCNPs maintained a net positive charge (∼+13 mV), which can be explained by the low charge shielding provided by HS15 compared to PEG-lipids with longer ethylene oxide segments, and the relatively high N/P ratio employed to load the LCNPs. To confirm the incorporation of siRNA into LCNPs, electrophoretic mobility shift assays (EMSA) were conducted at different N/P ratios. Gel images revealed that the siRNA cargo is mostly loaded in the particles as it remained trapped in the wells (fig. S1A). RiboGreen assays confirmed that at least 90% of siRNA was incorporated in the LCNPs at all tested N/P ratios, except the N/P ratio of 6 where it was lower (∼83%) (fig. S1B). It was also confirmed with EMSA that the siRNA was shielded from RNase-mediated degradation when incorporated in LCNPs, while naked siRNA or siRNA released from LCNPs was not (fig. S1C). In addition, no changes in the d_h_ and polydispersity index (PDI) of siRNA-loaded LCNPs were observed over a week (fig. S1D).

**Fig. 2.**
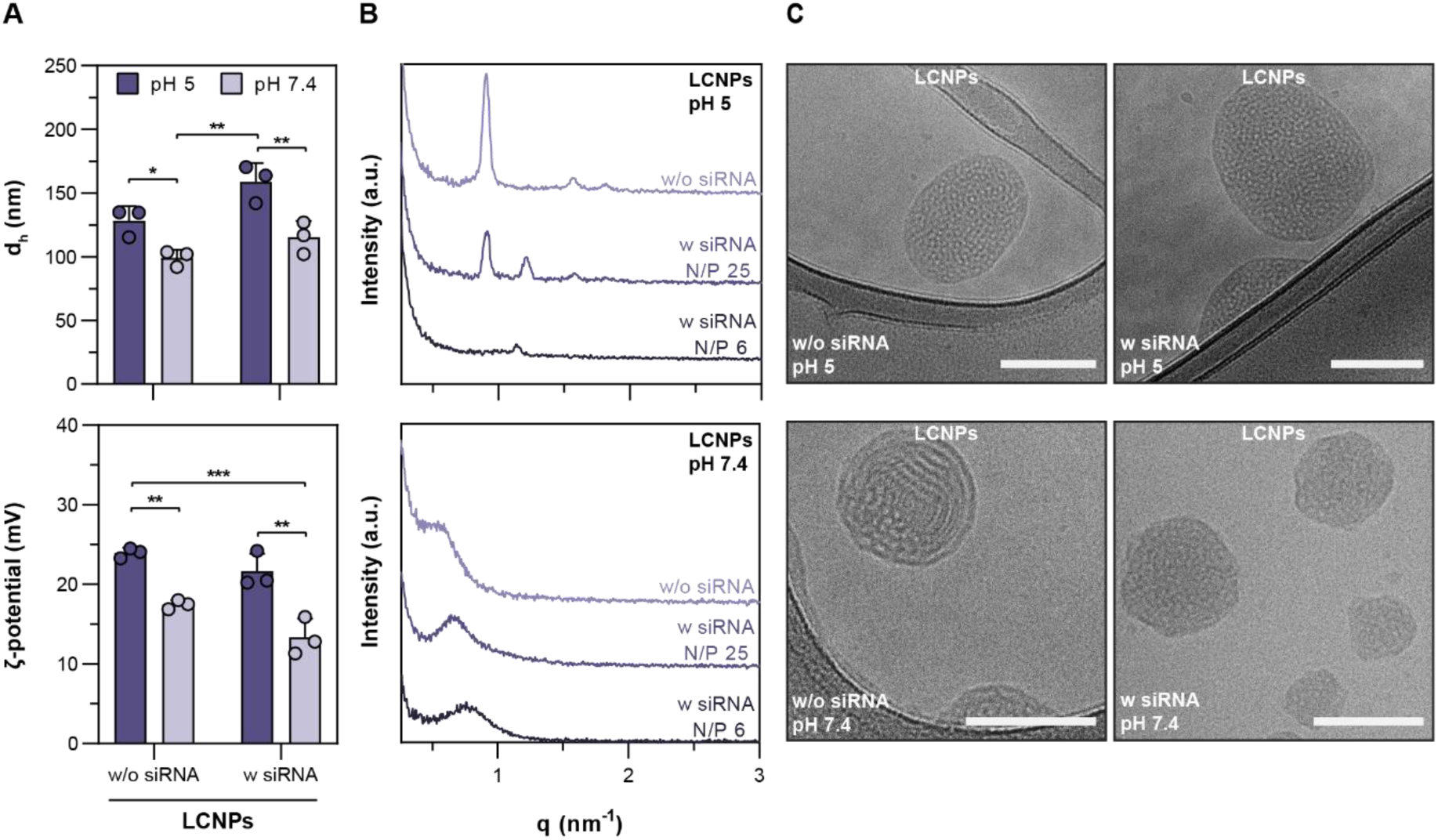
Physicochemical properties of LCNPs. LCNPs with a lipid/surfactant composition of MC3/DOPE/HS15 (60/18/22 mol%) exhibit pH-dependent structural changes. (**A**) d_h_ and ζ-potential of LCNPs at pH 5 and 7.4 with (N/P 25) or without siRNA (n = 3). (**B**) SAXS scans of LCNPs as a function of siRNA concentration (N/P ratio) and buffer pH (5 and 7.4). (**C**) Representative cryo-TEM images of LCNPs at pH 5 and 7.4 with (N/P 25) or without siRNA; scale bars 100 nm. Data are expressed as mean + s.d. of n independent experiments. Statistical significance was calculated from an ordinary two-way ANOVA with Tukey’s post-hoc test; *p < 0.05, **p < 0.01 and ***p < 0.001.

Next, we investigated the structural properties of LCNPs with SAXS. LCNPs exhibited a pH-dependent structure-switching behavior (Fig. 2B). SAXS scans of unloaded LCNPs at pH 5 displayed the formation of a well-defined H_II_ phase with a q-position ratio of 1:√3:2. The characteristic repeat distance (*d*) and the center-to-center distance (α) were estimated to be 6.93 nm and 8.01 nm, respectively. Upon introducing siRNA at an N/P ratio of 25, structural disorder increased, as indicated by reduced peak intensities, which eventually vanished at higher siRNA concentrations (N/P 6). Further, we observed the presence of an additional peak at a q value of 1.2 nm^-1^ for N/P 25 and N/P 6, suggesting the formation of a second population with smaller characteristic spacing (Fig. 2B). At pH 7.4, the higher-order peaks disappeared but the first structure peak persisted, whether or not siRNA was present. This observation suggests that a sponge (L_3_) phase formed at pH 7.4 (Fig. 2B). Interestingly, similar results were obtained recently for LNP cores, mixtures of MC3 (50 mol%) and cholesterol (38.5 mol%) (*47*). The peak position of the first structure peak was determined with a Lorentzian broad peak model, and it was observed that the peaks shifted to higher q values the more siRNA was added (fig. S2). In addition, the structure of LCNPs loaded with mRNA (N/P 25) was analyzed using SAXS, considering that the size and molecular structure of the loaded NA cargo could influence the nanostructural properties of LCNPs. A broad peak at low q (0.75 nm^-1^) suggested that L_3_ phases were preserved in mRNA-loaded LCNPs at pH 7.4 (fig. S3, refer to Supplementary Materials for a detailed discussion on physicochemical properties of these LCNPs). Cryo-TEM images of LCNPs corroborated the SAXS analysis. At both tested buffer pH values, LCNPs exhibited an electron-dense core in the presence or absence of siRNA (Fig. 2C). In contrast to conventional siRNA-loaded LNPs (*48, 49*), we observed circular patterns of non-lamellar phases at pH 5, whether with or without siRNA, and suggests the potential formation of H_II_ phases (Fig. 2C). Notably, these patterns were more disordered at pH 7.4 particularly in the presence of loaded siRNA, and LCNPs exhibited irregular spherical shapes similar to other L_3_-based nanoparticles (NPs) (Fig. 2C) (*50, 51*).

In addition, we investigated whether a structural transformation occurred when cone-shaped lipids were substituted with cylindrical lipids. Therefore, DOPE was replaced by DOPC or MC3 by DOTMA. A summary of the physicochemical properties of these DOPC- and DOTMA-based NPs are provided in fig. S4A. Like LCNPs, DOPC-NPs showed different structures at pH 5 and 7.4, which can potentially be attributed to the altered charge state of MC3 and stronger interactions with the NA cargo at acidic pH. Structure peaks in SAXS plots of DOPC-NPs suggested that multivesicular liposomes formed at pH 7.4. These structures became more irregular when loaded with siRNA (fig. S4B, and corresponding discussion in Supplementary Materials). Cryo-TEM images of unloaded DOPC-NPs at pH 7.4 clearly showed the formation of raspberry-shaped multivesicular systems (fig. S4C). Upon adding siRNA at pH 7.4, multi-vesicular/lamellar domains were found adjunct to domains with non-lamellar arrangements (fig. S4C). At pH 5, SAXS analysis revealed that DOPC-NPs exhibited hexagonal phases characterized by three sharp peaks with a q-position ratio of 1:√3:2, particularly when siRNA was added to the system (fig. S4B). Cryo-TEM images showed that DOPC-NPs formed well-defined non-lamellar structures at pH 5 (fig. S4C). As for DOTMA-NPs, we observed a series of broad oscillations with SAXS, suggesting bilayer formation with a low degree of multilamellarity both at pH 5 and 7.4 with or without siRNA (fig. S4D, and corresponding discussion in Supplementary Materials). Multi- and unilamellar liposomes smaller than LCNPs and DOPC-NPs (< 100 nm) were observed by cryo-TEM (fig. S4E). In line with the SAXS data, the structure of DOTMA-NPs did not change with respect to buffer pH variations and the presence of siRNA cargo.

### Characterization of EVs and hybridization kinetics

First, MSC-derived EVs were isolated from 3D cell cultures as previously reported by our group and characterized following the guidelines of the International Society for Extracellular Vesicles (ISEV) (*52–54*). In brief, MSC-EVs had a mean modal diameter of approximately 127 nm and a mean d_h_ of approximately 160 nm as determined by nanoparticle tracking analysis (NTA) and DLS, respectively. The ζ-potential was found to be around -28 mV (fig. S5A). The vesicular structure was confirmed by negative-stain transmission electron microscopy (TEM) and cryo-TEM (fig. S5B). Western blotting revealed the enrichment of EV-specific membrane markers, such as CD63 and CD9, in the vesicle isolates while cellular impurities like calregulin and Grp94 were absent (fig. S5C). Moreover, the MSC-specific membrane proteins CD73 and CD26 (DPP4) were highly enriched within EVs (fig. S5C).

Subsequently, we investigated the kinetics of HEV formation by monitoring particle concentration and d_h_ over a 60-min time course in a 384-well microplate reader setup using static light scattering (SLS) and DLS, respectively. We hypothesized that the hybridization of LCNPs with EVs should result in a decrease in the overall particle count and an increase in the overall size. When different number ratios of EVs were added to LCNPs (*i.e.*, 4:1, 1:1 and 1:4), particle concentrations decayed exponentially, stabilizing after approximately 30 min (fig. S6A). In contrast, the d_h_ increased over 30 min and plateaued at approximately 190 nm at all tested ratios (fig. S6A). Furthermore, we explored the impact of temperature and buffer pH (fig. S6, B and C) on HEV formation kinetics at an EV:LCNP number ratio of 1:1. As expected, the kinetics slowed down when the temperature was reduced from 37 to 4 °C, while no significant differences were noted in d_h_ under the tested temperature conditions (fig. S6B). With respect to pH, no differences were observed in the hybridization kinetics when HEVs were formed under basic conditions (pH 9) compared to pH 7.4 (fig. S6C). In contrast, as the pH was lowered (≤ 6), d_h_ dramatically increased reaching 350 nm at pH 5, and the particle concentrations could not be assessed anymore after the first 4 min as multimodal particle populations were detected, suggesting the formation of large aggregates (fig. S6C).

To elucidate the structural properties of the HEV formation, cryo-TEM imaging was performed. EVs and LCNPs were combined and imaged after 5 and 30 min and 4 h of incubation (pH 7.4, 37°C). When examined separately, EVs had an electron-lucent aqueous core (highlighted in blue) delimited by a circular, electron-dense lipid bilayer and siRNA-loaded LCNPs had an electron-dense core (highlighted in orange) with an irregular surface and shape (Fig. 3A). Membrane proteins on EV lipid bilayers appeared as darker, electron-dense spots (white arrows). Upon mixing and grid preparation (∼5 min), hemi-fused vesicles appeared, suggesting that hybridization processes between EVs and LCNPs can be instantly triggered (Fig. 3B). EV membrane proteins were visible in hemi-fused HEVs (white arrows), excluding the possibility that the structures are blebs typically found in mRNA-loaded LNPs (*55, 56*). Additionally, cryogenic electron tomography (cryo-ET) was performed to show that the hemi-fused HEVs are not superpositions of EVs and LCNPs (movie S1 and S2). Hemi-fused HEVs remained visible after 30 min of incubation (Fig. 3B). After 4 h, only a few unfused EVs and hemi-fused HEVs were still visible (Fig. 3B). Instead, we also observed larger, possibly fully-fused electron-dense structures (> 200 nm). The surface-bound membrane proteins were still visible as small black dots on these structures (white arrows), suggesting that EV membrane proteins are located on HEV surfaces post-hybridization (Fig. 3B).

**Fig. 3.**
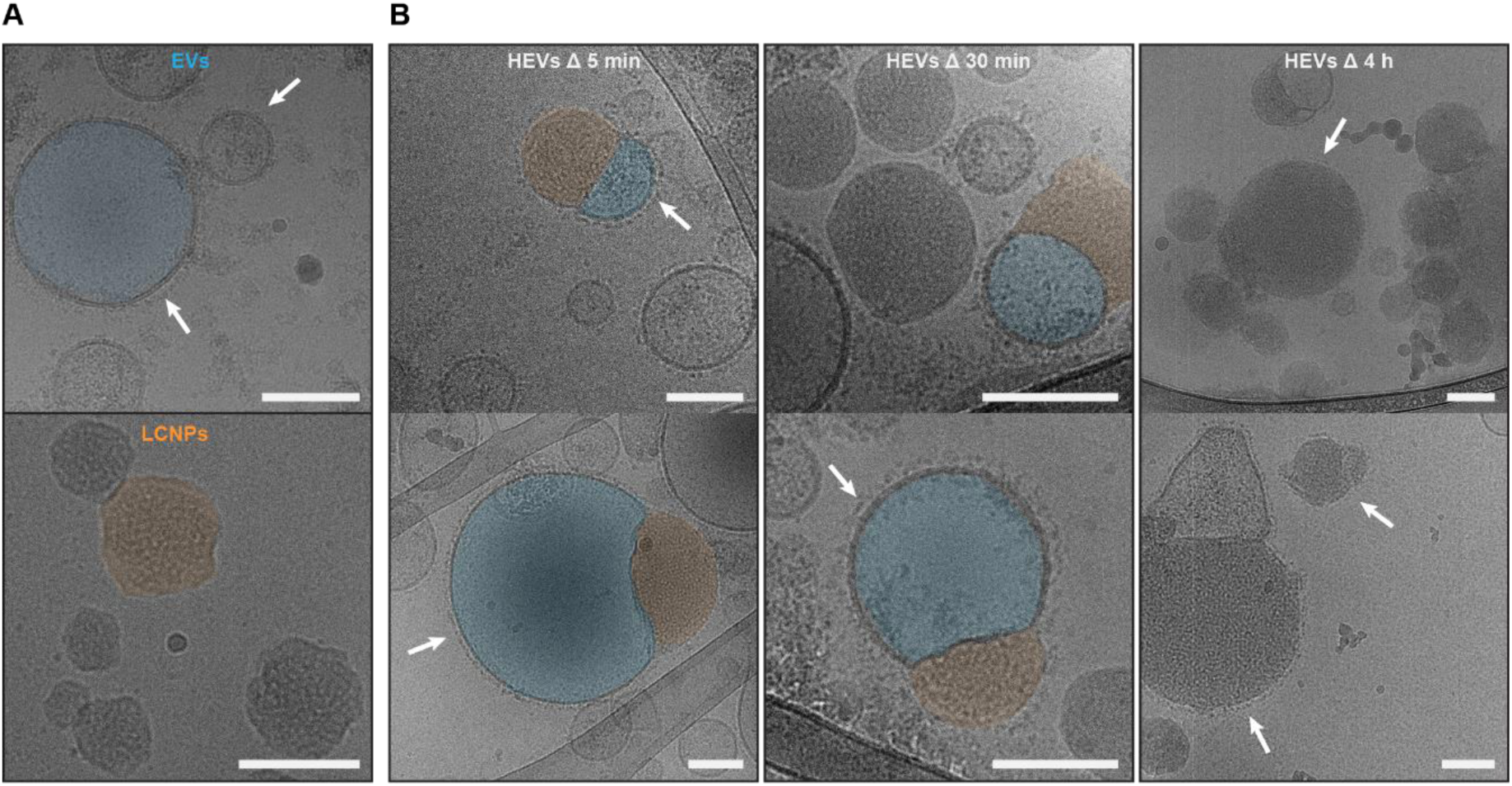
Cryo-TEM imaging of HEV formation. (**A**) Representative cryo-TEM images of MSC-EVs and siRNA-loaded LCNPs (N/P 25); scale bars 100 nm. EVs are characterized by an electron-lucent aqueous core (highlighted in blue) and LCNPs have a darker, electron-dense lipid core (highlighted in orange). EV membrane proteins are visible as darker, electron-dense spots (white arrows). (**B**) Time-dependent cryo-TEM images of mixtures of EVs and siRNA-loaded LCNPs (N/P 25) at a particle number ratio of 1:4 at 37 °C and pH 7.4; scale bars 100 nm. After 5 and 30 min, hemi-fused HEVs are visible (LCNP part highlighted in orange and EV part highlighted in blue). The presence of membrane proteins (white arrows) on the lipid bilayers of EVs rules out the possibility that the hemi-fused particles are phase separation phenomena within LCNPs. After 4 h of incubation mostly fully hybridized HEVs are visible. EV membrane proteins appear as dark electron-dense spots on the surface of HEVs (white arrows).

The presence of larger aggregates became apparent in cryo-TEM when EVs were mixed with LCNPs at pH 5, in line with our DLS results (figs. S7A and S6C). Similarly, DOTMA-NPs induced the formation of larger aggregate structures, whereas DOPC-NPs appeared to have minimal interaction with EVs as visualized by cryo-TEM (fig. S7A). However, owing to the more electron-lucent nature of DOPC-NPs compared to LCNPs, the possibility of hybrid formation or interaction with the EV membranes could not be ruled out. The formation of hemi-fused HEVs could also be observed when mRNA-loaded LCNPs and EVs were coincubated (fig. S7B).

### Topology of lipid nanocarriers influences their interactions with EV membranes

Nano-flow cytometry (NanoFCM) measurements were then conducted to characterize the HEV formation at the single-particle level. For this approach, EVs were labeled using CellTrace^TM^ Far Red, and combined with LCNPs, and DOPC-, as well as DOTMA-NPs loaded with the fluorescent FAM-siRNA at a particle number ratio of 1:1. The commercial Exo-Fect^TM^ system was used as a benchmark for comparison. As part of the negative controls, unlabeled EVs were mixed with the various NP formulations containing non-fluorescent siRNA. Additionally, labeled EVs were co-incubated with an equal quantity of naked FAM-siRNA.

Bivariate plots of single-stained EVs and NPs clearly showed distinct single-positive populations for CellTrace^TM^ Far Red (highlighted by the red square inset) and FAM-siRNA (highlighted by the dark green square inset), respectively (Fig. 4A). We observed that not all NPs were loaded with fluorescent siRNA (*e.g.*, for LCNPs at pH 7.4 ∼38%) (fig. S8). This may stem from rearrangements in the lipid matrix during the pH adjustment step or the relatively high N/P ratio of 25, resulting in loaded and unloaded LCNPs (*57*). Notably, for the Exo-Fect^TM^ transfection reagent no FAM-siRNA positive events were detected. As reported previously, this discrepancy may stem from the relatively low particle concentration found in Exo-Fect^TM^-siRNA-transfection-mixtures, which in our case was three orders of magnitude lower compared to our NPs (*e.g.*, 4.9 × 10^10^ particles/mL for Exo-Fect^TM^ *vs.* 1.5 × 10^13^ particles/mL for LCNPs) (*39*). When different FAM-siRNA-loaded NP formulations were mixed with CellTrace^TM^-labeled EVs, a double-positive population appeared, indicating the formation of HEVs (highlighted by the lime green square inset) (Fig. 4A). This double-positive population was not present when, for example, naked FAM-siRNA was coincubated with labeled EVs or in the case of the Exo-Fect^TM^ transfection reagent (Fig. 4A).

**Fig. 4.**
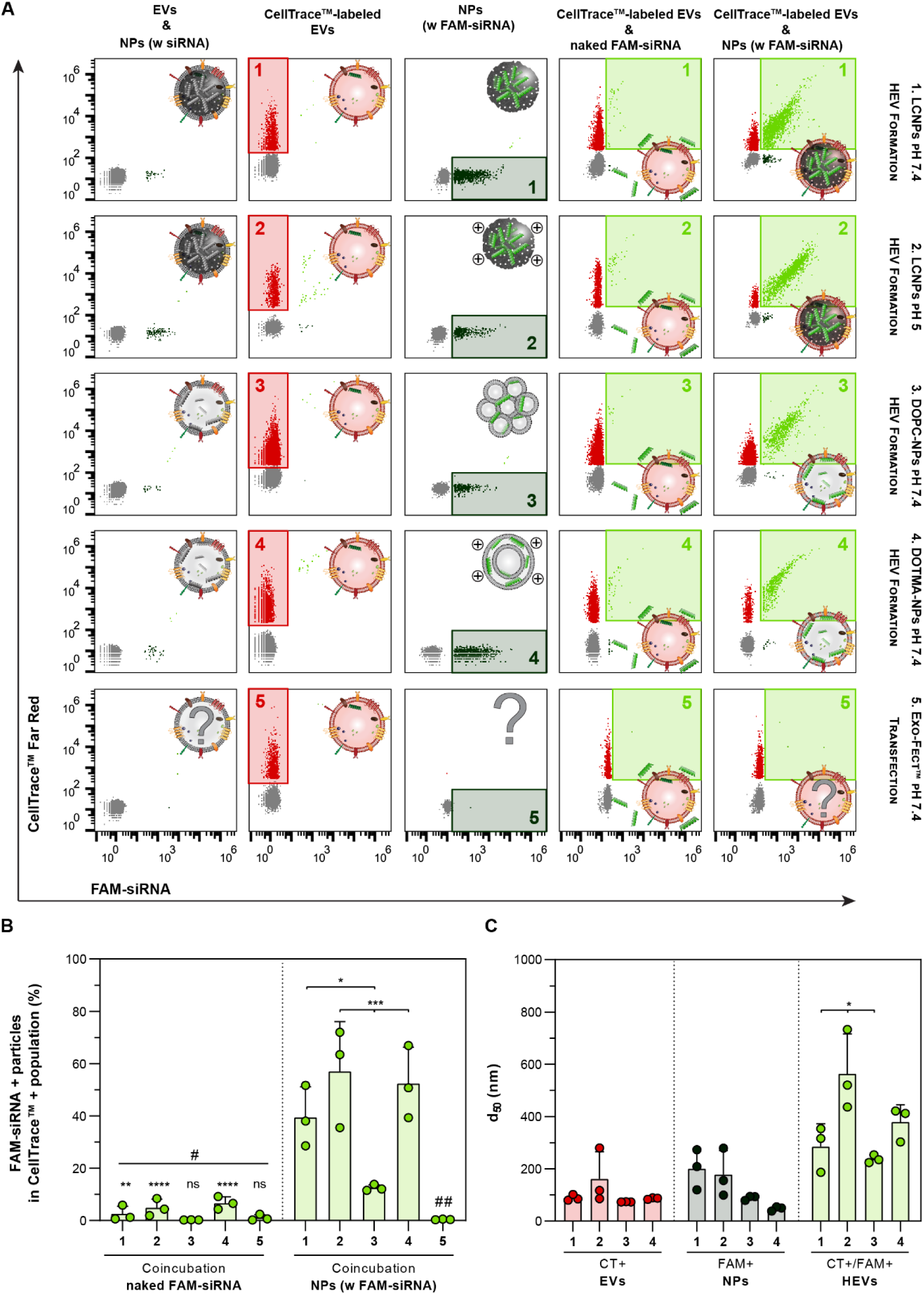
Topology and physicochemical properties of lipid nanocarriers affect HEV formation. CellTrace^TM^ Far Red-labeled EVs are combined at a particle number ratio of 1:1 with the different FAM-siRNA-loaded (N/P 25) lipidic NPs (LCNPs, DOPC- and DOTMA-NPs) or with the Exo-Fect^TM^ reagent (∼2 μM FAM-siRNA; 10 min at 37 °C). After 30 min of incubation at 37 °C the formation of HEVs is assessed by NanoFCM. (**A**) Representative bivariate dot plots of FAM-siRNA+ particles (x-axis) *vs.* CellTrace^TM^+ particles (y-axis). Particle species are labeled from left to right: mixtures of unlabeled EVs and NPs are used as labeling controls. CellTrace^TM^-labeled EVs with the red square insets highlight the CellTrace^TM^+ particle populations. Different FAM-siRNA-loaded NPs with the dark green square insets highlight the FAM-siRNA+ particle populations. Coincubation of naked FAM-siRNA (∼2 μM) with CellTrace^TM^-labeled EVs with the lime green square insets highlight the double-positive (CellTrace^TM^+/FAM-siRNA+) particle populations. Coincubation of FAM-siRNA-loaded NPs with CellTrace^TM^-labeled EVs with the lime green square insets highlight the double-positive (CellTrace^TM^+/FAM-siRNA+) particle population. Hybridization conditions are labeled from 1 to 5: (1) LCNPs pH 7.4; (2) LCNPs pH 5; (3) DOPC-NPs pH 7.4; (4) DOTMA-NPs pH 7.4; (5) Exo-Fect^TM^ pH 7.4. (**B**) Percentages of the double-positive particles (CellTrace^TM^+/FAM-siRNA+) are shown relative to the respective total CellTrace^TM^+ particle populations after coincubation with naked FAM-siRNA, FAM-siRNA-loaded NPs or Exo-Fect^TM^ (n = 3). (**C**) d_50_ of the individual particle populations: CellTrace^TM^+ EVs; FAM-siRNA+ NPs; and CellTrace^TM^+/FAM-siRNA+ HEVs (n = 3). Data is expressed as mean + s.d. of n independent experiments. Statistical significance was calculated from an ordinary one-way ANOVA with Tukey’s post-hoc test; *p < 0.05, **p < 0.01, ***p < 0.001, ****p < 0.0001, ns = not significant; # = *vs.* respective experimental condition; ## = *vs.* 1 ***p < 0.001, *vs.* 2 & 4 ****p < 0.0001.

Comparing the percentages of double-positive particle counts relative to the total single-positive CellTrace^TM^ population, a clear trend could be observed (Fig. 4B). HEV formation with highly positively charged NPs, like LCNPs under acidic conditions or DOTMA-NPs, resulted in the highest colocalizations of approximately 57 and 53%, respectively. The interaction between LCNPs and EVs decreased to 40% at pH 7.4 and to 12% when multi-vesicular/lamellar DOPC-NPs were used instead (Fig. 4B). This suggests that surface charge, lipid compositions and thus the NP topology may influence the interaction of NPs with EV membranes. When EVs were coincubated with free FAM-siRNA or with the Exo-Fect^TM^ reagent, the double-positive population was at background levels below 7% (Fig. 4B).

The size profiles of individual particle populations were also determined with NanoFCM. Single-positive EV and FAM-siRNA-loaded NP populations had median diameters (d_50_) ranging from 50 to 200 nm (Fig. 4C). In the double-positive HEV populations, we observed a size shift towards larger particles. The d_50_ increased to 290 and 240 nm upon mixing EVs with LCNPs or DOPC-NPs at pH 7.4, respectively. At pH 5, a drastic size shift to 560 nm was observed when EVs were incubated with LCNPs and the d_50_ reached 380 nm when DOTMA-NPs were used instead (Fig. 4C). This observation aligns well with DLS and cryo-TEM results, confirming that highly positively charged NPs form larger structures potentially induced by aggregation with EVs (figs. S6C and S7A).

To further confirm the mixing of LCNP lipids with EVs, 1 mol% of DOPE was substituted with Atto 488-labeled DOPE. Also in this case, a double-positive population appeared when EVs were combined with LCNPs (highlighted by the lime green square inset) (fig. S9A). This double-positive population accounted for 45% of the total CellTrace^TM^+ population at pH 7.4 and 75% at pH 5 (fig. S9B). Moreover, the size of HEVs increased from 170 to 490 nm when the pH decreased from 7.4 to 5 in line with our previous observations (fig. S9C).

To validate the LCNP platform for EV loading, we also evaluated HEV formation using different NA cargos, including mRNA and an RNA aptamer. Cy5-labeled mRNA was loaded into LCNPs and combined with carboxyfluorescein succinimidyl ester (CFSE)-stained EVs. Similar to the siRNA experiments, a double-positive population appeared, constituting 41% of the total CFSE-positive population (highlighted by the pink square inset) (fig. S9D). Furthermore, we utilized the RNA aptamer Broccoli for the *in situ* detection of HEV formation (*58, 59*). The membrane-permeable dye DFHBI-1T diffuses into both HEVs and LCNPs and specifically binds to the Broccoli aptamer. The formation of Broccoli-loaded HEVs was only detected when CellTrace^TM^-labeled EVs were mixed with Broccoli-loaded LCNPs in presence of the DFHBI-1T dye (highlighted by the lime green square inset), amounting to 15% of the total CellTrace^TM^+ population (fig. S9E). The detection of double-positive events may have been affected by dilution of the freely diffusing dye inside the flow cell, potentially leading to a reduced detection rate compared to experiments with covalently labeled NAs.

### Detection of EV membrane proteins on HEV surfaces using an optofluidic platform

To investigate whether EV membrane proteins are present on HEV surfaces, and whether particular subpopulations of EVs preferentially interact with LCNPs, we applied a state-of-the-art optofluidic platform based on an integrated microfluidic chip that combines immunoaffinity pull-down assays with correlative single molecule interferometric scattering and fluorescence microscopy (fig. S10, see details in Supplementary Materials). The microfluidic chip integrated all steps of the immunoassay, encompassing surface functionalization using supported lipid bilayers, antibody conjugation, and EV/HEV sensing. Conceptually this platform isolates EV subpopulations on the sensor substrate by immobilizing EVs expressing specific surface markers, and then distinguishes between HEVs loaded with fluorescent NA cargo and native EVs by correlating the scattered and fluorescence signals from each pulled-down vesicle (Fig. 5A). For correlative label-free and fluorescence mapping, we sequentially imaged the EV/HEV mixtures on the same camera by alternating between off- and on-resonant fluorescence excitations, respectively. Particles visible in both imaging modalities in the presence of specific EV-markers were assigned as NA-loaded HEVs. LCNPs were immobilized on the surface of the sensor, *via* their higher non-specific binding to the functionalized substrate compared to EVs and HEVs. This difference in non-specific binding between particle species can be attributed to residual electrostatic interactions between the negatively charged groups of the functionalized surface, caused by the biotin and neutravidin moieties, with the net positive ζ-potential of LCNPs (Fig. 2A).

**Fig. 5.**
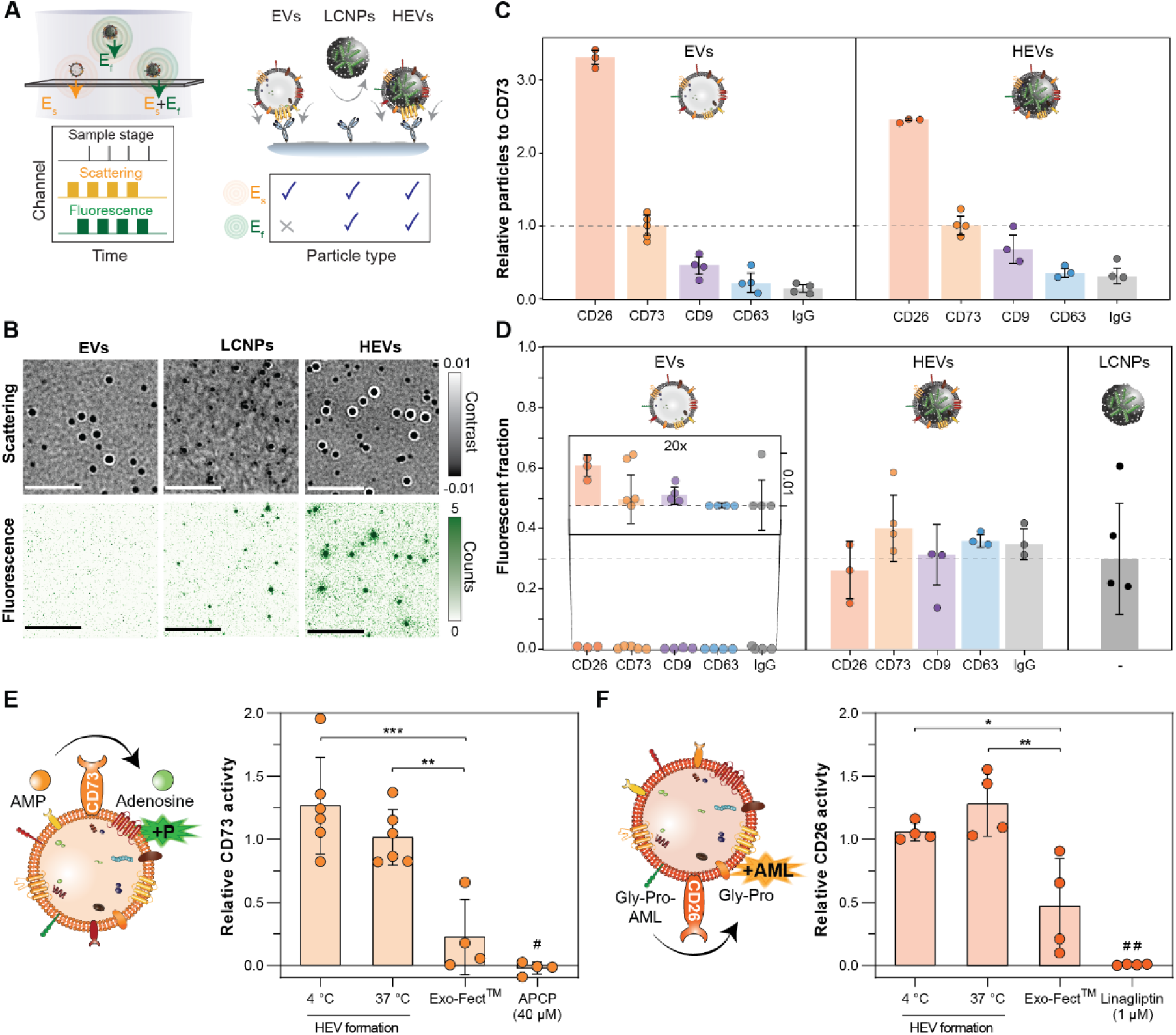
In-depth characterization of HEVs: surface protein expression, colocalizaton with fluorescent NA cargo, and enzymatic activity post-hybridization. Native MSC- EVs are mixed at a particle number ratio of 1:4 with FAM-siRNA-loaded (N/P 25) LCNPs and after 30 min of incubation at 37 °C particle populations are analyzed with an optofluidic platform. (**A**) Conceptual schematic of the correlative label-free and fluorescence optofluidic platform used to simultaneously determine the FAM-siRNA loading of each particle population and surface protein expression profile. Left: synchronization scheme between the channels controlling sample scanning, label free excitation/detection and fluorescence excitation/detection. Right: particle assignment table resulting from the pull- down immunoaffinity assay. (**B**) Representative images of each particle type under the two detection schemes; scale bars 5 μm. (**C**) Relative EV-specific surface protein expression levels for EVs and HEVs normalized to the number of CD73+-pulled down particles (n = 3-5). (**D**) Fluorescent fraction resulting from correlative scattering and fluorescence signals for EVs and HEVs expressing a particular surface protein, and for non-specifically bound LCNPs (n = 3-5). Data is represented as the median over n independent experiments ± s.d. (**E**) A malachite green assay was employed to determine the activity of the EV membrane protein CD73 after HEV formation (1:1 particle number ratio; 30 min at 4 and 37 °C), treatment with the Exo-Fect^TM^ reagent (10 min at 37 °C) and the CD73 inhibitor (40 μM APCP) (n = 4-6). Enzyme activity was normalized to EVs subjected to identical hybridization conditions. (**F**) A protease assay was employed to assess the activity of the EV membrane protein CD26 after HEV formation (1:1 particle number ratio; 30 min at 4 and 37 °C), treatment with the Exo-Fect^TM^ reagent (10 min at 37 °C) and the CD26 inhibitor (1 μM linagliptin) (n = 4). Enzyme activity was normalized to EVs subjected to identical hybridization conditions. Data is expressed as mean ± s.d. of n independent experiments. Statistical significance was calculated from an ordinary one-way ANOVA with Tukey’s post-hoc test; *p < 0.05, **p < 0.01, ***p < 0.001; # = *vs.* HEV (37 °C) ***p < 0.001, *vs.* HEV (4 °C) ****p < 0.0001; ## = *vs.* HEV (37 °C) ****p < 0.0001, *vs.* HEV (4 °C) ***p < 0.001.

Fig. 5B shows representative zoom-in images (12 µm × 12 µm) of the three different particle types under the two different imaging modalities. As expected, EVs exhibited scattering signals but no fluorescent ones. LCNPs showed weaker scattering signals compared to the EVs, but also displayed a fraction of coincident fluorescent signatures. This can be attributed to the fact that not all LCNPs were loaded with FAM-siRNA, as demonstrated by NanoFCM (fig. S8). HEVs exhibited stronger scattering signals compared to EVs that also coincided with a fraction of fluorescent ones, indicating the successful hybridization of EVs with LCNPs (fig. S11).

To establish the molecular signature of HEVs and compare it to their native EV source, we selected a panel of antibodies targeting various EV- and MSC-specific membrane proteins as capture probes. Multiplexing of the immunoaffinity assays was achieved by functionalizing spatially separated sensing channels with different antibodies targeting the known EV markers CD9 and CD63, along with MSC-specific surface markers CD73 and CD26. To determine the intrinsic level of non-specific binding for each particle species, we used IgG1 as an isotype control. For each sensing channel replica, we imaged a total surface area of 0.1 mm^2^ and analyzed between 10^3^-10^4^ particles per biomarker. To obtain the relative expression levels of EVs/HEVs positively identified with each surface marker, we normalized the total EVs and HEVs with respect to the MSC-specific protein CD73. For EV samples, we observed the following trend in the relative expression profile of membrane proteins CD26 >> CD73 > CD9 > CD63 > IgG, quantified in terms of the number of pulled down particles (Fig. 5C). Notably, the expression profile did not change after HEV formation, generated with a 4:1 number excess of LCNPs over EVs. This suggests that the membrane proteins are located on the HEV surfaces in agreement with our cryo-TEM observations (Figs. 5C and 3B).

Taking advantage of the multiplexing capabilities of the microfluidic-based platform, we could determine the association of the FAM-siRNA cargo with HEVs for several specific EV subpopulations within a single experiment at single-particle resolution. Therefore, the number of double-positive events from superimposed fluorescent and label-free interferometric scattering signals were quantified *via* image analysis. The number of colocalizations fell within the range of 30 to 40% for all tetraspanin markers, with no statistically significant difference between them (Fig. 5D). This suggests that LCNPs do not preferentially hybridize with any specific EV subpopulation. In agreement with the NanoFCM data, control experiments with LCNPs utilizing non-specific electrostatic interactions for immobilization revealed that only 30% were loaded with the fluorescent siRNA (Fig. 5D). Since only 30% of LCNPs contained FAM-siRNA cargo, and the HEVs were prepared with a 4:1 excess of LCNPs to EVs for hybridization, we infer that nearly all the fluorescent NA cargo was transferred to EVs.

### Biological activity of membrane proteins and cargo integrity is retained post-hybridization

Since we could confirm that MSC-specific membrane proteins CD73 and CD26 are located on the surface of HEV populations, we measured their enzymatic activity post-hybridization using a colorimetric and a luminescence-based plate assay, respectively. The CD73 activity was not affected after HEV formation. In contrast, it decreased significantly when EVs were treated with the Exo-Fect^TM^ reagent or the inhibitor control (40 μM APCP) (Fig. 5E). This confirms previous results of altered EV membrane integrity after treatment with Exo-Fect^TM^ (*39*). Likewise, we did not observe a reduction of the CD26 enzymatic activity, while the Exo-Fect^TM^ reagent and the inhibitor control (1 μM linagliptin) impaired it (Fig. 5F).

To evaluate the integrity of exogenously loaded NA cargo in HEVs and endogenous NA cargo in EVs, we employed two different strategies. First, HEVs loaded with the Broccoli aptamer were co-incubated with RNase A at 37 °C for 24 h, and the fluorescence signal was tracked in the presence of the aptamer-selective dye DFHBI-1T. In case the aptamer is released from HEVs, it would be susceptible to RNase-mediated cleavage, resulting in reduced interaction with DFHBI-1T, and a subsequent decline of the fluorescence signal over time (fig. S12A). In Broccoli-loaded HEVs the fluorescence signal remained stable over 24 h both in the presence and absence of RNase. Similarly, Broccoli-loaded LCNPs maintained their fluorescence signal in both conditions. Conversely, the naked Broccoli displayed a complete loss of fluorescence signal in the presence of RNase (fig. S12B). In addition, an automated gel electrophoresis instrument was employed to analyze the integrity of endogenous intraluminal NA cargo encapsulated in EVs before and after HEV formation. Therefore, EVs were hybridized with empty LCNPs, treated with RNase, and total RNA was extracted for analysis. Distinct RNA bands appeared in the electropherogram of both EV and HEV samples (fig. S12C). A slight shift and a reduction in the fluorescence intensities of the RNA peaks were visible in HEVs compared to the EV sample, suggesting that a fraction of the natural RNA cargo might get lost during the hybridization process (fig. S12D). However, additional quantitative analyses are needed to validate these observations.

### HEVs are taken up by cells but do not enhance endosomal membrane disruption

Before performing uptake studies, cytotoxicity tests with LCNPs at concentrations ranging from 12.5 to 200 µg/mL for 4 or 24 h, both in the presence and absence of encapsulated siRNA were conducted. Compared to the Triton^TM^ X-100 positive control, no significant changes in cell viability were observed for all tested conditions (fig. S13). Next, we investigated whether LCNPs and HEVs could efficiently enter cells and deliver their NA cargo. We therefore labeled LCNPs both with the fluorescent Cy5-siRNA and the fluorescent lipid Atto 488 DOPE. Following hybridization with MSC-EVs the dual-labeled HEVs and LCNPs were incubated with HeLa cells for 4 and 24 h and the uptake was analyzed with flow cytometry. After 4 h, the median fluorescence intensities (MFI) of Cy5 and Atto 488 signals significantly increased compared to the PBS control, confirming the successful uptake of LCNPs and HEVs (Fig. 6A). Confocal images of HeLa cells treated with Cy5-siRNA-loaded HEVs or LCNPs corroborated the flow cytometry results, displaying bright, punctate Cy5 signals both within cells treated with LCNPs and HEVs (fig. S14). After 24 h LCNPs exhibited 2-fold higher MFI values compared to HEVs (Fig. 6A). Similar results were previously reported for EV-liposome hybrids (*32*).

**Fig. 6.**
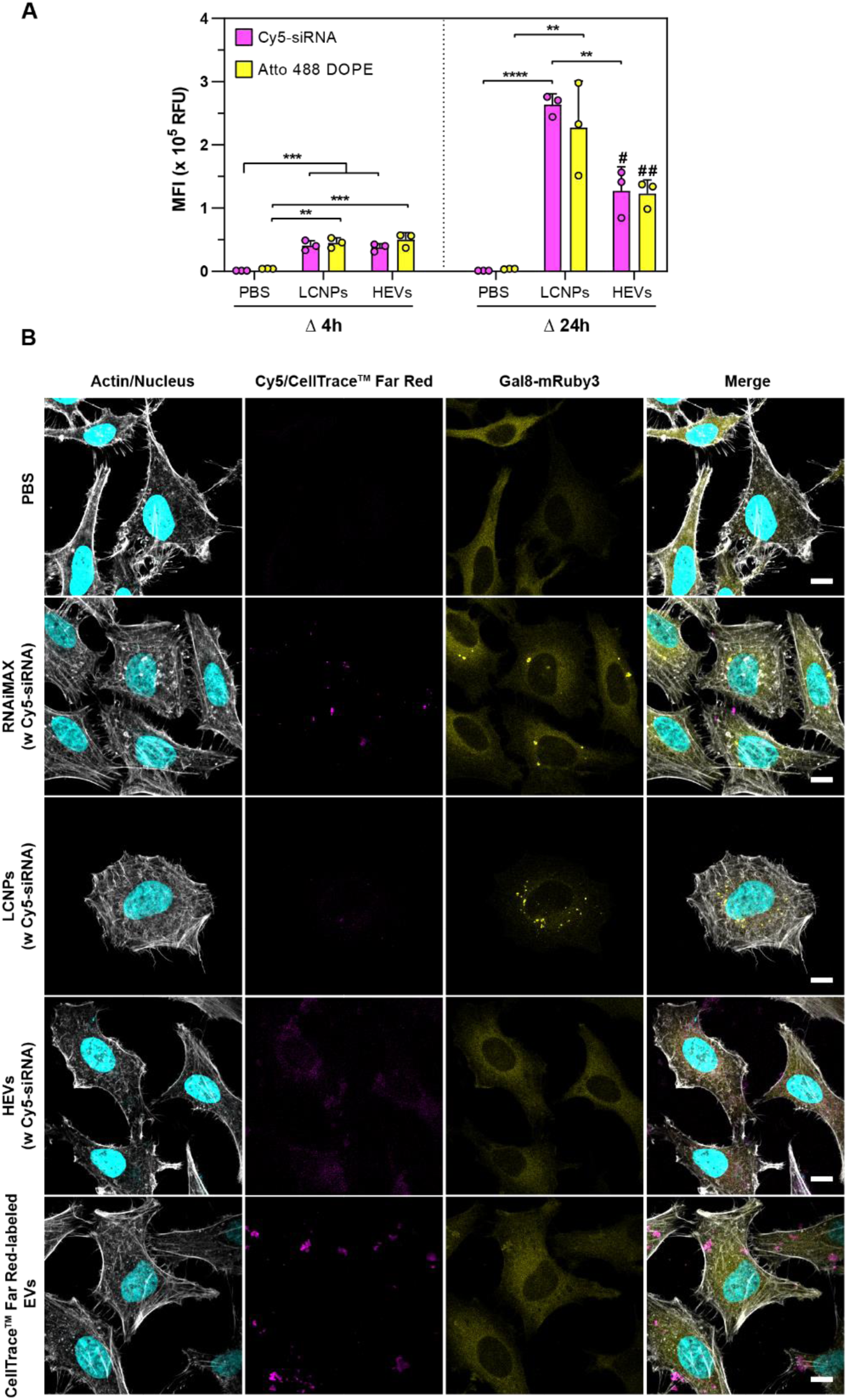
Cell uptake and endosomal escape of HEVs and LCNPs. (**A**) Uptake of Cy5- siRNA (N/P 25)/Atto 488 DOPE (1 mol%) dual-labeled HEVs and LCNPs in HeLa cells after 4 and 24 h of incubation (n = 3). HEVs were prepared by incubating EVs and LCNPs at a 1:1 particle number ratio for 30 min at 37 °C. Cells were treated with 20 nM Cy5- siRNA/well. (**B**) Endosomal escape assay with HeLa cells stably expressing the Galectin-8 mRuby3 fusion protein (Gal8-mRuby3). Cytosolic Gal8-mRuby3 interacts with β- galactoside sugars exposed upon endosomal damage and results in bright punctate signals. The panel shows the four different imaging channels, from left to right: actin/nucleus (gray/cyan), Cy5/CellTrace^TM^ Far Red (magenta), Gal8-mRuby3 (yellow) and the merged image. HeLa-Gal8-mRuby3 cells were treated for 4 h with Cy5-siRNA-loaded LCNPs (N/P 25) and HEVs at a concentration of 100 nM Cy5-siRNA/well. Lipofectamine^TM^ RNAiMAX with Cy5-siRNA (100 nM Cy5-siRNA/well) and PBS served as positive and negative controls, respectively. CellTrace^TM^ Far Red-labeled EVs (∼3.4 × 10^10^ particles/mL) were added for comparison. Scale bars are 10 μm. Data is expressed as mean + s.d. of n independent experiments. Statistical significance was calculated from an ordinary one-way ANOVA with Tukey’s post-hoc test; **p < 0.01, ***p < 0.001 and ****p < 0.0001; # = *vs.* PBS **p < 0.01; ## = *vs.* PBS *p < 0.05.

In addition, we investigated the endosomal remodeling in HeLa cells stably expressing the Galectin-8 mRuby3 fusion protein (Gal8-mRuby3). NP-mediated disruption of the endosomal membrane causes the recruitment and binding of Gal8 to β-galactoside sugars located on the inner leaflet of endosomes (*60, 61*). The binding of Gal8-mRuby3 can then be followed by a transition from a diffuse cytosolic signal to bright punctate patterns. Therefore, HeLa-Gal8-mRuby3 cells were imaged 4 h after addition of Cy5-siRNA-loaded HEVs and LCNPs. Native MSC-EVs labeled with CellTrace^TM^ Far Red and Lipofectamine^TM^ RNAiMAX were employed as controls. Both native EVs and Lipofectamine^TM^ carriers were taken up by cells as visualized from the signals of intracellular CellTrace^TM^ and Cy5-siRNA signals (Fig. 6B). In alignment with previous studies, EVs did not induce endosomal damage, in contrast to the Lipofectamine^TM^ positive control, where a clear enhancement of the punctate mRuby3 signal could be observed (Fig. 6B) (*62, 63*). Notably, HEVs displayed no endosomolytic effects whereas LCNPs induced bright mRuby3 punctate signals (Fig. 6B), suggesting potential differences in their interactions with endosomal membranes and/or different uptake pathways. Further detailed mechanistic investigations are warranted to comprehensively understand the intracellular trafficking of HEVs and LCNPs.

### HEVs show improved knockdown and expression efficiencies *in vitro*

The knockdown efficiency of HEV and control formulations was assessed in a HeLa cell line stably expressing GFP. Specifically, HeLa-GFP cells were subjected to treatment with siGFP-loaded HEVs and LCNPs at a 10 nM siRNA concentration (5 pmol per well). Lipofectamine^TM^ RNAiMAX was employed as positive control. Negative controls consisted of HEVs and LCNPs loaded with a scrambled control siRNA (siCTRL), as well as naked siGFP and PBS. The GFP expression levels were assessed 48 h post-sample addition by flow cytometry. Additionally, representative fluorescence microscopy images are provided in fig. S15.

Under low serum conditions (8%, v/v), siGFP-loaded HEVs reduced the GFP fluorescence intensity by approximately 75%, akin to the Lipofectamine^TM^ RNAiMAX positive control (Fig. 7A). Similarly, LCNPs led to a significant reduction of the GFP fluorescence by approximately 60% (Fig. 7A). LCNPs loaded with the siCTRL did not affect the GFP expression, similar to cells treated with naked siGFP or PBS. In contrast, HEVs loaded with the siCTRL elicited a modest reduction in GFP expression by approximately 20% (Fig. 7A). This might be due to the components of MSC-derived EVs that may interact with the innate immune system (*e.g.*, through toll-like receptors (TLRs)) and/or deliver gene-modulating NAs (*e.g.*, miRNAs) and thereby momentarily decrease GFP expression levels (*64, 65*).

**Fig. 7.**
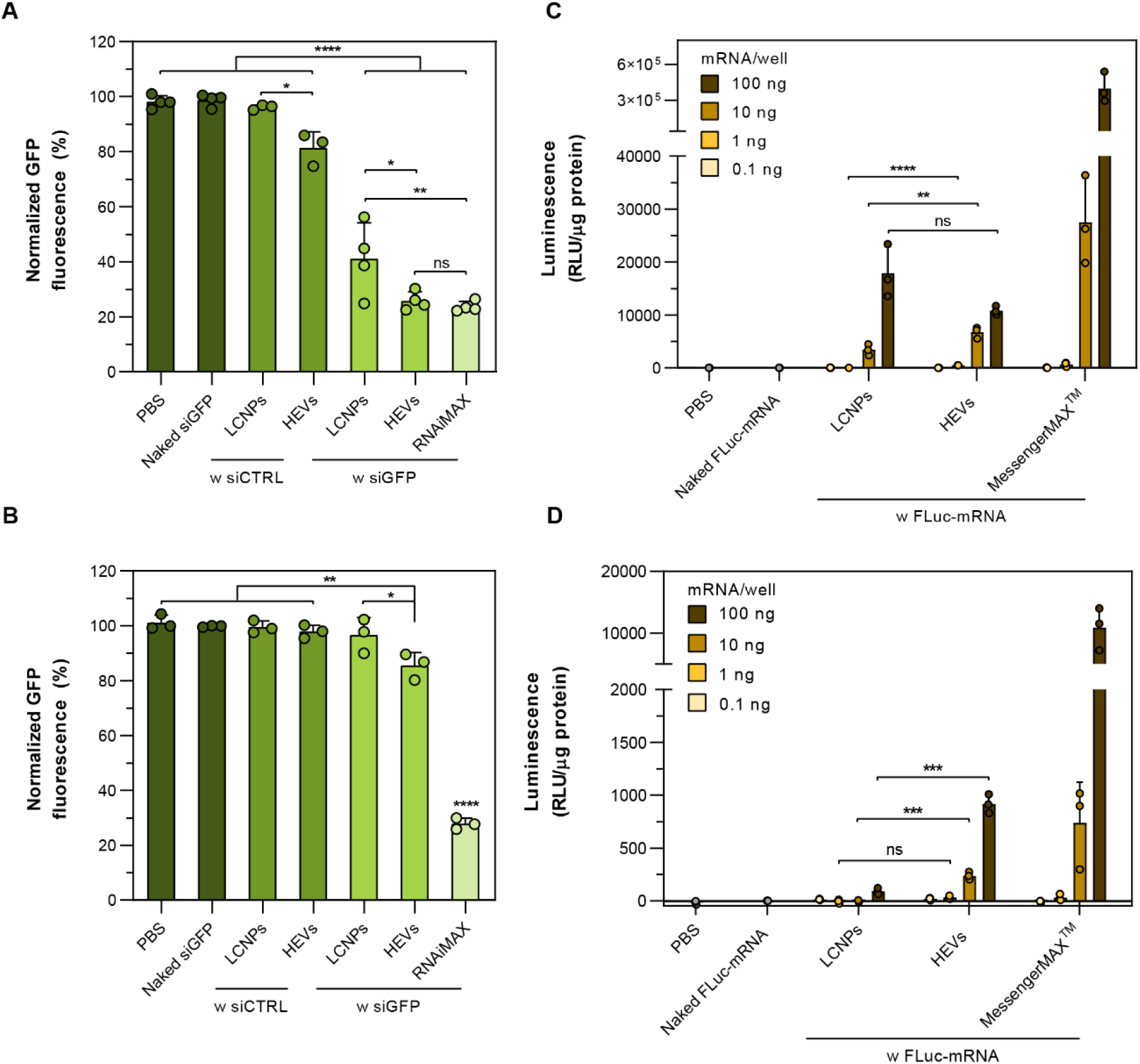
*In vitro* knockdown and expression efficiencies of HEVs. HEVs were prepared by incubating EVs with siRNA- or mRNA-loaded LCNPs (both N/P 25) for 30 min at 37 °C and particle number ratios of 1:1. (**A**) GFP knockdown in 8% (v/v) serum after 48 h of incubation with siRNA-loaded LCNPs and HEVs (10 nM/well). GFP fluorescence is analyzed 48 h after sample addition; siCTRL is a scrambled siRNA not targeting the GFP mRNA (n = 3-4). (**B**) GFP knockdown in 80% (v/v) serum after 4 h of incubation with siRNA-loaded LCNPs and HEVs (10 nM/well). GFP fluorescence is analyzed 48 h after sample addition (n = 3). (**C**) mRNA dose-dependent expression of the firefly luciferase (FLuc) in 10% (v/v) or (**D**) in 90% (v/v) serum after 24 h of incubation with FLuc-mRNA- loaded LCNPs and HEVs (n = 3). Data is expressed as mean + s.d. of n independent experiments. Statistical significance was calculated from an ordinary one-way ANOVA with Tukey’s post-hoc test in (**A**) and (**B**). Statistical differences between LCNP and HEV transfections in (**C**) and (**D**) were assessed with a two-tailed unpaired Student’s t-test instead of ANOVA tests, given the pronounced expression profile of the Lipofectamine^TM^ MessengerMAX^TM^ positive control; *p < 0.05, **p < 0.01, ***p < 0.001, ****p < 0.0001 and ns = not significant.

Subsequently, identical experiments were conducted under more challenging cell culture conditions, reducing the contact time with cells from 48 to 4 h, and increasing the serum concentration to 80% (v/v). HEVs retained some transfection capability, leading to a 15% reduction in GFP fluorescence compared to the untreated control (Fig. 7B). In contrast, LCNPs loaded with siGFP, along with formulations containing siCTRL or naked siGFP, all exhibited GFP expression levels equivalent to those of untreated cells (Fig. 7B). Notably, the gold standard Lipofectamine^TM^ RNAiMAX maintained a robust knockdown efficiency even under high serum conditions (Fig. 7B).

We then assessed firefly luciferase (FLuc) expression in HeLa cells transfected with FLuc-mRNA-loaded HEVs or LCNPs at mRNA doses ranging from 0.1 to 100 ng/well (0.3 pM to 0.3 nM). Expression levels were quantified by measuring bioluminescence signals after 24 h of incubation. Under low serum conditions (9%, v/v), both mRNA-loaded LCNPs and HEVs exhibited similar expression levels at a high mRNA dose of 100 ng/well (Fig. 7C). With decreasing the mRNA dose to 10 and 1 ng/well, HEVs showed significantly higher signal intensities compared to LCNPs (Fig. 7C). As expected, the gold standard Lipofectamine^TM^ MessengerMax^TM^ demonstrated the highest transfection efficiencies across all tested mRNA doses. At 0.1 ng/well of mRNA, all formulations exhibited bioluminescence signals comparable to the negative controls (naked FLuc-mRNA and PBS). Under high serum conditions (90%, v/v), we observed a significant increase in expression levels with HEVs in comparison to LCNPs at mRNA doses of 100 and 10 ng/well (Fig. 7D). In summary, the *in vitro* transfection results suggest that HEVs demonstrate superior transfection efficiency compared to LCNPs, especially in high serum conditions and at low mRNA doses. Similar observations were also recently reported in a comparative study of *in vitro* transfection experiments with sgRNA-loaded EVs and LNPs (*18*).

## Discussion

Recent structural investigations on LNPs have shown that the structural organization of the lipid matrix can be modulated by the process parameters (*e.g.*, buffer pH and automated *vs.* microfluidic preparation protocols) and the lipid composition (*66–68*). It was shown that LNPs with non-lamellar LLC domains *(i.e.*, H_II_ phases or the bicontinuous cubic phases (Q_II_)) display improved transfections *in vitro* and *in vivo* (*69, 70*). This was partially attributed to their enhanced fusogenic properties leading to increased endosomal escape and cytosolic NA delivery. In this work, we investigated the relationship between non-lamellar LLC phases in LNPs and their hybridization with EVs. A ternary mixture of the ionizable lipid MC3, the helper lipid DOPE and the non-ionic surfactant HS15 resulted in the non-lamellar LCNPs with a pH-dependent structure-switching behavior from H_II_ phase at pH 5 to L_3_ phase at pH 7.4 (Fig. 2). Highly positively charged H_II_-based LCNPs aggregated with EVs when incubated at pH 5. In contrast, a controlled hybridization process could be achieved under physiological conditions with L_3_ phase-structured LCNPs. Interestingly, the multi-vesicular/lamellar DOPC-based NPs interacted less with EVs, whereas multilamellar NPs formed with the permanently positively charged lipid DOTMA led to the formation of aggregates (Figs. 3 and 4). This suggests that there is a defined interplay between membrane surface potential and nanostructural properties of lipid nanocarriers and their successful hybridization with EVs. At this point, we can only hypothesize why L_3_-based LCNPs promote fusion processes with EV membranes and further mechanistic studies are needed to better understand the mechanism. In a recent study by Valldeperas et al. the interfacial behavior of lipidic NPs with L_3_ phases were studied (*51*). It was shown that upon interaction with hydrophilic surfaces L_3_-based NPs spread, collapsed and rearranged into thin lipid layers. Since EVs are mixtures of endosome- and plasma membrane-derived lipid vesicles, their negative surface potential and the presence of anionic lipids, such as phosphatidylserine, could in a similar way induce structural rearrangements within L_3_-based LCNPs (*71, 72*). Furthermore, recent findings suggest that particularly DOPE interacts strongly with EV membranes, highlighting its important role in the hybridization process (*73*). NanoFCM revealed that various NA molecules (*i.e.*, mRNA, siRNA and RNA aptamer) could be incorporated into EVs, highlighting the versatility of the LCNP platform for loading EVs. The optofluidic platform presented in this study enabled us to study EV/HEV subpopulations based on the expression of membrane proteins (Fig. 5). It revealed that the membrane proteins of MSC-EVs and the fluorescent NA cargo were transferred onto/into HEVs, and around 30 to 40% of HEV particles contained the fluorescent NA cargo irrespective of the EV subpopulation. Importantly, we confirmed that the intrinsic biological activity of MSC-EV enzymes CD73 and CD26 was retained post-hybridization. Finally, *in vitro* transfection results highlighted the enhanced transfection capabilities of HEVs compared to their hybridization partner LCNPs, while minimally perturbing the endosomal membrane (Figs. 6 and 7). Future studies should aim at further investigating the *in vivo* fate and therapeutic potential of HEVs. In particular, whether they will be avidly sequestered by the mononuclear phagocyte system or, depending on the EV source, exhibit organotropism (*74*).

## Materials and Methods

### Materials

The human bone marrow stromal cell line HS-5 and HeLa cells were acquired from ATCC (USA). HeLa-GFP cells were kindly provided by Dr. Tom Edwardson (Department of Chemistry and Applied Biosciences, ETH Zurich, Switzerland) and were initially purchased from Cell Biolabs, Inc. (USA). HeLa-Gal8-mRuby3 cell line was a kind gift from Dr. Simone Berger (Department of Pharmacy, LMU Munich, Germany). The pET28c-F30-Broccoli plasmid was a kind gift from Dr. Samie Jaffrey (Department of Pharmacology, Cornell University, USA) (Addgene plasmid # 66788). Dulbecco’s modified eagle medium (DMEM), phenol red-free DMEM, fetal bovine serum (FBS), GlutaMAX^TM^ supplement, penicillin-streptomycin (10,000 U/mL), minimal essential medium (MEM) non-essential amino acids (NEAA) solution (100x), Opti-MEM^TM^, trypsin-EDTA (0.25%), UltraPure^TM^ DNase/RNase-free distilled water, UltraPure^TM^ agarose, phosphate buffered saline (PBS), Dulbecco’s phosphate buffered saline (DPBS), HEPES 1 M solution, ethanol absolute, Hoechst 33342 and blasticidin S hydrochloride (from *Streptomyces Griseochromogenes*) were purchased from Thermo Fisher Scientific (USA). Sodium chloride (NaCl), sodium acetate, sodium acetate solution 3 M, uranyl acetate, sodium fluoride (NaF), sodium orthovanadate (Na_3_VO_4_), sodium dodecyl sulfate (SDS), sodium hydroxide (NaOH), potassium chloride (KCl), ammonium persulfate, phenylmethanesulfonyl fluoride (PMSF), boric acid, tris(hydroxymethyl)aminomethane (Tris) base, ethylenediaminetetraacetic acid (EDTA), urea, Triton^TM^ X-100, Tween^®^ 20, guanosine-5’-monophosphate (GMP), β-mercaptoethanol, isopropanol, tetramethylethylenediamine (TEMED), adenosine 5’-(α,β-methylene)diphosphate (APCP), methanol-free paraformaldehyde (PFA), skim milk powder, bovine serum albumin (BSA) and RNase A from bovine pancreas were purchased from Sigma-Aldrich (USA). Kolliphor^®^ HS 15 was obtained from BASF (Germany). Magnesium chloride (MgCl_2_) and ammonium molybdate tetrahydrate was purchased from abcr GmbH (Germany). Bromphenol blue was purchased from Alfa Aeser (USA). Malachite green was from Bender & Hobein GmbH (Switzerland). Adenosine-5’-monophosphate (AMP) was purchased from Acros Organics (Belgium). Glycerol, hydrochloric acid (HCl) 37% and acetic acid glacial were obtained from VWR Chemicals (USA). Tris/acetate/EDTA (TAE) buffer 50x was purchased from AppliChem GmbH (Germany). cOmplete^TM^ EDTA-free protease inhibitor cocktail was from Roche (Switzerland). Acrylamide bis-acrylamide 30% solution, 29:1 was from Bio-Rad (USA). DFHBI-1T was purchased from Tocris Bioscience (UK). DLin-MC3-DMA was purchased from MedChemExpress (USA). DOPE and linagliptin were purchased from AdipoGen Life Sciences (USA). DOPC was purchased from Fluorochem (UK). Atto 488 DOPE was purchased from ATTO-TEC GmbH (Germany). DOTMA was purchased from Larodan AB (Sweden). Detailed product-specific information of the various NA sequences and antibodies used in this study are found in tables S1 and S2.

### *In vitro* transcription of Broccoli RNA aptamer

The Broccoli RNA aptamer was synthesized by *in vitro* transcription. Initially, DNA templates were generated from the pET28c-F30-Broccoli template plasmid using PCR and utilizing the Phusion^®^ high-fidelity DNA polymerase (NEB, UK), forward primer 5’-ATA CCC ACG CCG AAA CAA- 3’ and reverse primer 5’-_M_G_M_GG AGC CCA CAC TCT ACT-3’ (where _M_G represents 2’-O-methoxyethyl (MOE)-modified guanine). The PCR products were subsequently purified using the QIAquick PCR Purification Kit (Qiagen, Germany). *In vitro* transcription was performed overnight at 37 °C with the MEGAshortscript^TM^ T7 Transcription Kit (Thermo Fisher Scientific) adding 5 mM GMP and 4 U/µL RNaseOUT^TM^ recombinant ribonuclease inhibitor (Thermo Fisher Scientific) to the transcription reaction*. In vitro*-transcribed RNA was treated with TURBO^TM^ DNase (Thermo Fisher Scientific) and purified using an 8% denaturating polyacrylamide gel prepared in Tris/borate/EDTA (TBE) buffer (100 mM Tris, 90 mM boric acid and 1 mM EDTA) containing 8 M urea. Electrophoresis was conducted at an electric field strength of 15 V/cm, and RNA bands were visualized using UV shadowing. RNA bands were excised from the gel, crushed with a pipet tip and the RNA was extracted overnight at 4 °C in water containing 0.3 M sodium acetate. The gel extracts were filtered through glass wool and the RNA was purified by isopropanol precipitation and dissolved in water. The quality and purity of the Broccoli RNA aptamer were confirmed by measuring A260/A280 and A260/A230 ratios, with both ratios approximating 2.0. Additionally, Broccoli RNA aptamers were analyzed with liquid chromatography mass spectrometry (LC-MS) (Supplementary Materials) and detailed sequence-specific information and physical properties can be found in table S1.

### Cell culture

The bone marrow-derived mesenchymal stem cell line HS-5, HeLa, and HeLa-Gal-8-mRuby3 (*60, 75*) cells were cultured in DMEM supplemented with 10% (v/v) FBS, 100 U/mL penicillin and 100 µg/mL streptomycin. HeLa-GFP cells were cultured in phenol red-free DMEM supplemented with 0.1 mM MEM NEAA, 2 mM GlutaMAX^TM^ supplement, 100 U/mL penicillin, 100 µg/mL streptomycin and 10 µg/mL blasticidin. To maintain the quality of cell cultures, subculturing was performed with low passage numbers (< 20), and routine mycoplasma contamination checks were conducted using the MycoAlert^TM^ Mycoplasma Detection Kit (Lonza, Switzerland).

### 3D culture of HS-5 cells

EV production from HS-5 cells was conducted using a 3D culture method previously described (*53*). In brief, 118 × 10^6^ HS-5 cells were seeded onto 4.5 g of Corning^®^ enhanced attachment microcarriers (Corning, USA) in a total volume of 300 mL full medium inside 500 mL Corning^®^ reusable glass spinner flasks (Corning). To promote cell attachment, an intermittent stirring protocol was applied overnight (5 min on at 30 rpm and 60 min off). Subsequently, the impeller speed was set to 35 rpm. After 48 h, the microcarriers were washed twice with PBS and 400 mL serum- and phenol red-free medium was added to the spinner flasks. The culture supernatant was collected after an additional 48 h of culture in serum-free medium.

### EV isolation from cell culture supernatant

EVs from cell culture supernatant were isolated according to previously published protocols (*52, 53*). Briefly, the supernatant was first processed by centrifugation at 500 and 2000 × *g* for 5 min each and 10,000 × *g* for 20 min using a Sorvall^TM^ LYNX 6000 centrifuge equipped with a Fiberlite^TM^ F12-6 × 500 LEX fixed-angle rotor (both Thermo Fisher Scientific). All centrifugation steps were performed at 4 °C. The clarified cell culture supernatant was then filtered through a 0.2-µm membrane before proceeding with ultracentrifugation (UC). UC was performed using an Optima XE-90 centrifuge (Beckman Coulter, USA) and a Type 45 Ti fixed-angle titanium rotor. After the first UC run (70 min at 4 °C and 100,000 × *g*), EV pellets were washed with PBS or HEPES-buffered saline (HEBS) (20 mM HEPES, 150 mM NaCl, pH 7.4) and pooled together. A second UC run was performed with identical conditions. EVs were suspended in appropriate volumes of ice-cold PBS/HEBS and stored in high concentrations (∼1 × 10^12^ particles/mL) at -20 °C.

### Preparation of NA-loaded LCNPs

NA-loaded LCNPs and formulation variations were produced using a microfluidic micromixer. Specifically, DLin-MC3-DMA or DOTMA, along with helper lipids (DOPE or DOPC), and the surfactant Kolliphor^®^ HS 15, were mixed at a molar ratio of 60:18:22 (cationic ionizable lipid/helper lipid/surfactant) in ethanol to achieve a final concentration of 20 mM (∼12 mg/mL). NAs (siRNA, mRNA, and RNA aptamer) were dissolved in 125 mM sodium acetate buffer at pH 4 (*76, 77*). Detailed sequence-specific information and physical properties of the various NAs employed in this study can be found in table S1. For particles containing fluorescent lipids, 1 mol% of DOPE was replaced by Atto 488-labeled DOPE. A staggered microfluidic diffusion mixing chip with a Y-shaped sample inlet (Fluidic 186, Microfluidic ChipShop, Germany) and a dual syringe pump system (Nemesys^®^, CETONI GmbH, Germany) were employed to combine the organic and aqueous phases at a volumetric flow rate ratio of 1.5:1 (buffer to ethanol) and a total flow rate of 0.05 mL/min. Following mixing, the ethanol concentration reached 40% (v/v), which was subsequently removed by dialysis against a 1000-fold volume excess of PBS (pH 7.4), HEBS (pH 7.4), or Broccoli aptamer buffer (20 mM HEPES, 150 mM KCl, 1 mM MgCl_2_, pH 7.4) for a minimum of 4 h using 20-kDa molecular weight cut-off dialysis units (Thermo Fisher Scientific). For pH variation experiments the sample buffer was adjusted with 62.5 mM HCl or NaOH solution and monitored with an InLab Micro^TM^ pH electrode (Mettler Toledo, USA). Unless otherwise specified, all formulations containing NAs were prepared with N/P ratios of 25, corresponding to a siRNA concentration of approximately 0.15 mg/mL and an mRNA/aptamer concentration of roughly 0.16 mg/mL in the final formulations.

### Western blotting

EV-specific proteins (CD73, CD26, CD63, CD9 and TSG101) as well as contamination markers (Grp94 and calregulin) were detected *via* western blotting. To obtain EV lysates, EV suspensions were diluted 5:1 (v/v) with 5x lysis buffer (comprising 100 mM Tris, 750 mM NaCl, 25 mM EDTA, 5% (v/v) Triton^TM^ X-100, 125 mM NaF, 5 mM PMSF, 5 mM Na_3_VO_4_, and 5x cOmplete^TM^ EDTA-free protease inhibitor cocktail). For cell lysates, cell pellets (approximately 0.2 × 10^6^ cells) were incubated with 100 µL of 1x lysis buffer. Protein concentration was determined using the Micro BCA^TM^ Protein Assay Kit (Thermo Fisher Scientific) following the manufacturer’s instructions. Lysates were combined 5:1 (v/v) with 5x Laemmli sample buffer (containing 0.25 M Tris, 10% (w/v) SDS, 30% (v/v) glycerol, and 0.02% (w/v) bromophenol blue), with or without reducing agent (5% (v/v) β-mercaptoethanol), boiled at 95 °C for 10 min and centrifuged at 10,000 × *g* for 1 min. Protein samples (5-10 µg) were resolved on a 12% SDS polyacrylamide gel at 80 V for approximately 1.5 h. The transfer to polyvinylidene difluoride (PVDF) membranes was performed with a Trans-Blot^®^ Turbo^TM^ system (at 25 V for 10 min, Bio-Rad). Blocking was conducted for 2 h using Tris-buffered saline (20 mM Tris, 150 mM NaCl) with 0.1% (v/v) Tween^®^ 20 (TBS-T) and 5% (w/v) skim milk powder. Primary antibodies were incubated overnight at 4 °C. Following extensive TBS-T washing, secondary horseradish peroxidase (HRP)-conjugated antibodies in TBS-T were added for 2 h at room temperature. Antibody concentrations are provided in table S2. Blots were developed by incubating them with western blotting luminol reagent (Santa Cruz Biotechnology, USA) for 2 min and immediately imaged using a ChemiDoc MP instrument (Bio-Rad).

### Size and ζ-potential measurements

Particle concentrations and size distributions of the various nanocarriers were determined with NTA using a ZetaView^®^ PMX 120-Z instrument (Particle Metrix GmbH, Germany), featuring a 405-nm laser and a COMOS camera. The measurements were conducted at 25 °C with sensitivity and shutter settings at 85 and 150, respectively. Videos were recorded at 11 different positions, each with two readings, and a frame rate of 30/s. To ensure accurate measurements, samples were diluted in PBS to achieve a concentration of at least 100 events per visual field. ζ-potential measurements were carried out with the same acquisition settings in the pulsed mode.

The d_h_, PDI and ζ-potential was also assessed with DLS on a Malvern Zetasizer Advance Pro instrument (Malvern Panalytical, UK) operating at a scattering angle of 179° and at a temperature of 25 °C. Samples were diluted 1:10 (v/v) with PBS at pH 7.4 or 5 before each measurement. The ζ-potential was determined *via* laser Doppler anemometry using the same instrument. Samples were diluted 1:50 (v/v) with 1 mM KCl solution at pH 7.4 or 5 before each measurement.

### SAXS analysis

For SAXS measurements, a sample volume of 20 μL was added to borosilicate glass capillary tubes (length 80 mm and wall thickness 0.01 mm, Hilgenberg GmbH, Germany) using hypodermic stainless-steel needles. Subsequently, the capillaries were hermetically sealed using 2-K-epoxy resin adhesive (resin to hardener ratio of 1:1 (v/v), UHU^®^ Plus). Capillaries were inserted into a specialized sample holder and measurements were performed using an in-house SAXS setup (Xeuss 3.0) equipped with a Genix 3D source (both Xenocs, France). The X-ray energy was 8.05 kV (λ = 1.54 Å, CuKα), and the photon flux at the sample position was > 10^7^ counts/s. The beam size on the sample was approximately 0.5 × 0.5 mm^2^ FWHM. The sample-to-detector distance, calibrated using a silver behenate standard, was 500 mm and the scattered signal was collected on a Dectris Eiger 1M detector (1028 × 1062 pixels, pixel size 75 × 75 µm^2^, Dectris Ltd., Switzerland). In total, the signal was recorded for > 8400 s. SAXS data are shown in the q-range from 0.2 to 6.9 nm^-1^, corresponding to real length scales from 1.05 to 39.25 nm.

Detector data were masked, azimuthally integrated, normalized to absolute intensity (cm^-1^) and background subtracted using XSACT 2.6 software package (Xenocs). The integrated intensity I(q) is plotted against the magnitude of the scattering vector q calculated with Eq. 1

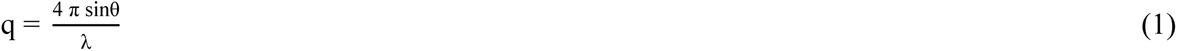

where *θ* is the scattering angle and *λ* is the wavelength of the incident X-rays. In case of indexing of sharp peaks, this procedure was carried out on the I(q) profiles before subtraction of the buffer background data, which appeared flat in the relevant q-range, to reduce the appearance of noise. Model intensities for the broad peak models were calculated using the open-source scattering analysis software SasView 5.0.5 (https://www.sasview.org/).

### EMSA

For EMSA experiments, 0.1 µg of naked siRNA or siRNA encapsulated within LCNPs were mixed with PBS to reach a final volume of 40 µL and 10 µL of a 70% (v/v) glycerol solution in PBS was added. The mixtures were then loaded onto a 0.5% (w/v) agarose gel and run for 30 min at 70 V in 0.5x TAE buffer (20 mM Tris-base, 10 mM acetic acid, and 0.5 mM EDTA). Subsequently, the gel was stained for 45 min with a 1:10,000 (v/v) dilution of SYBR^TM^ Gold Nucleic Acid Gel Stain (Thermo Fischer Scientific) in 0.5x TAE buffer and visualized using the ChemiDoc MP imaging system. To investigate the protection of the siRNA cargo from nuclease degradation, a similar experiment was conducted in the presence of RNase A. Specifically, 5 µg/mL RNase A was added to the samples, with or without 1% (v/v) Triton^TM^ X-100 and incubated for 1 h at 37 °C before loading onto gels.

### Determination of the loading efficiency

The loading efficiency (LE) of NAs within the various nanocarriers was determined with the Quant-it^TM^ RiboGreen RNA Assay Kit (Thermo Fisher Scientific) according to manufacturer’s instruction. Briefly, the mean fluorescence intensities of unecapsulated NAs (MFI_u_) were determined and subtracted from the total fluorescence intensities (MFI_t_) measured after lysing the nanocarriers with 1% Triton^TM^ X-100. Fluorescence measurements were carried out on a Tecan Spark^®^ plate reader (Tecan, Switzerland) with excitation set to 480 ± 10 nm and emission set to 520 ± 10 nm. A calibration curve from 20 ng/mL to 1 μg/mL was used to determine the absolute NA concentration and LE (in %) was calculated according to Eq. 2:

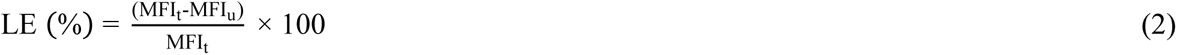

### Cell viability

Cell viability was assessed using the CellTiter 96^®^ AQueous One Solution Cell Proliferation (MTS) Assay (Promega, Switzerland). One day before treatment, 2,500 HeLa cells were seeded into 96-well plates and were incubated overnight at 37 °C and 5% CO_2_. Subsequently, the cells were treated with escalating concentrations of LCNPs (ranging from 12.5 to 200 μg/mL), either unloaded or loaded with siRNA (N/P ratio of 25). PBS and 1% (v/v) Triton^TM^ X-100 were used as controls. After 4 and 24 h of incubation, the cells were washed twice with PBS, and the medium was replaced with 100 μL of DMEM. Twenty μL of the MTS reagent was added to each well. After 1 h of incubation at 37 °C and 5% CO_2_, the absorbance was measured at 490 nm using the Tecan Spark^®^ plate reader. Cell viability (in %) was determined by comparing the treated conditions to an untreated control.

### Preparation of HEVs

HEV formation was consistently carried out using a freshly prepared LCNP suspensions. HEVs were prepared by combining specified particle number ratios (4:1, 1:1 and 1:4) of EVs and LCNPs as determined by NTA. The mixtures were thoroughly vortexed and incubated at different temperatures (4, 25 and 37 °C) and for different durations (ranging from 5 min to 4 h). In experiments involving pH variations, a 62.5 mM HCl or NaOH solution was used to amend the sample buffer pH and was readjusted to pH 7.4 before measurements. Unless specified otherwise, HEVs were formed at a pH of 7.4 and a temperature of 37 °C, with an EV:LCNP number ratio of 1:1, over a 30-min incubation time.

### DLS/SLS time-course experiments

Hybridization kinetics were evaluated through DLS/SLS time-course experiments, employing a DynaPro Plate Reader III (Wyatt Technology, USA). The experiments were performed under temperature-controlled conditions (4, 25, and 37 °C) within sealed 384-well microplates (Aurora Microplates, USA). Instrument settings were configured as follows: laser auto-attenuation was enabled, and each well was subjected to 4 acquisitions, each lasting 3 s. Measurements at 4 °C were performed under a nitrogen atmosphere. Before each measurement, the microplate was pre-incubated with 27 μL of a freshly prepared LCNP suspension (∼9 × 10^8^ particles/mL) for at least 30 min until a stable particle concentration reading was achieved. Subsequently, 10 µL of an equivalent particle amount (except for ratio variation experiments) of EVs was added, and the measurement (particle concentration and d_h_) was continued for 60 min. The total particle amount was kept constant at ∼4.9 × 10^7^ particles/well.

### Electron microscopy and tomography

For TEM imaging, 5 µL of EV suspensions in PBS (∼1 × 10^11^ particles/mL) were adsorbed onto glow-discharged 300-mesh copper grids. The grids underwent a 30-s glow discharge at 25 mA using a PELCO easiGlow^TM^ glow discharge cleaning system (Ted Pella, USA). Subsequently, the grids were stained with a 2% (w/v) uranyl acetate solution and blotted with filter paper. Finally, they were rinsed with distilled water, blotted again with filter paper, and dried under ambient conditions. EV samples were imaged with a Tecnai F20 field emission gun (FEG) microscope (Field Electron and Ion Company, USA) operating at a 200-kV acceleration voltage. The microscope was equipped with a combination of CCD (Gatan ORIUS SC200 2K, AMETEK Inc., USA) and direct electron detector (Falcon II 4K, Thermo Fisher Scientific).

For cryo-TEM imaging, 3.6 μL of HEV, LCNP or EV suspensions in PBS (∼1 × 10^12^ particles/mL) were deposited on 300-mesh lacey carbon-coated copper grids (Quantifoil Micro Tools GmbH, Germany). The grids were previously glow-discharged for 45 s using an Emitech K100X glow discharge system (Quorum Technologies Ltd., UK). Excess liquid was removed by blotting for 2 s at 100% relative humidity before immersion of the grids into a mixture of liquid ethane and propane using a Vitrobot Mark IV (Thermo Fisher Scientific). For the automated specimen loading procedure, the vitrified grids were clipped into AutoGrid specimen carriers (Thermo Fisher Scientific).

The prepared grids were then examined in bright-field mode using a TFS Titan Krios (Thermo Fisher Scientific) operated at 300 kV acceleration voltage and equipped with a Gatan Quantum LS Energy Filter (GIF) using a 20 eV slit width and a Gatan K2 Summit direct electron detector (AMETEK Inc.). The microscope was maintained at a temperature of approximately -180 °C during observations. Micrographs were acquired in an energy filtered TEM (EFTEM) operation mode using the TFS EPU software (up to 105,000x magnification, at most ∼40 e/Å^2^ total electron dose, K2 camera in a linear mode, within 2-4 μm defocus range). Images were processed using Fiji software (78).

Tomographic data collection was performed using the Tomo software on a Titan Krios G3i microscope (all Thermo Fisher Scientific) equipped with a Gatan BioQuantum imaging filter (GIF) with a slit width of 20 eV and K3 direct electron detector (AMETEK Inc.). The carbon-coated grids were prepared as described above. The tilt series was acquired using the dose-symmetric tilt scheme from -40° to +40° in 2° tilt steps at 19,500x magnification with a pixel size of 0.44 and a total electron dose of ∼130 e/Å^2^. The reconstructions and visualizations were performed using the software package Inspect3D (Thermo Fisher Scientific) by choosing the simultaneous algebraic reconstruction technique (SART) protocols implemented in the software.

### NanoFCM analysis

EVs were labeled using either a 5 µM solution of CellTrace^TM^ Far Red (Thermo Fisher Scientific) or 5(6)-carboxyfluorescein diacetate *N*-succinimidyl ester (CFDA-SE) (Chemodex AG, Switzerland) in PBS as previously described (*78*). In brief, EVs were mixed with the respective dyes and incubated overnight at 4 °C. Subsequently, the mixtures were incubated at 37 °C for 15 min to facilitate acetate hydrolysis of the dye. Unbound dye was removed using dialysis against a 1000-fold excess of PBS or Broccoli aptamer buffer at 4 °C overnight with the Pur-A-Lyzer^TM^ Mini 6000 Dialysis Kit (Sigma-Aldrich).

Different NPs, formulated either with fluorescently tagged NAs or lipids (FAM-siRNA, Cy5-mRNA, or Atto 488 DOPE) or with the Broccoli aptamer, were then mixed with fluorescently labeled EVs at a particle number ratio of 1:1 (∼7 × 10^11^ particles/mL). The formation of HEVs was evaluated using a Flow NanoAnalyzer N30 instrument (NanoFCM Co., Ltd., UK). For comparison, the Exo-Fect^TM^ Exosome Transfection Kit (System Biosciences, USA) was used following manufacturer’s instruction, with an equivalent amount of FAM-siRNA (∼2 μM). Control experiments were conducted by coincubating with an equivalent amount of naked fluorescent NAs (∼2 μM for FAM-siRNA, ∼0.1 μM for Cy5-mRNA and ∼6 μM for Broccoli aptamer), and using non-fluorescent EVs and NPs or by excluding the Broccoli aptamer selective dye DFHBI-1T from the Broccoli aptamer detection buffer (20 mM HEPES, 150 mM KCl, 1 mM MgCl_2_, and 20 µM DFHBI-1T).

Events were recorded for 1 min on three different channels: 488/10, 525/40 and 670/30 with a single-photon counting avalanche photodiode detector (SPCM APD). Channel settings were aligned using 250 nm fluorescent silica QC beads and size calibration curves were generated using the S17M-MV size standard (150-1000 nm) (all NanoFCM Co., Ltd.). For experiments involving FAM-siRNA, Atto 488 DOPE, and Broccoli aptamer, the blue 488-nm laser was set to 10/50 mW, and the red 638-nm laser was set to 40/100 mW, with a SS decay of 0.2%. For Cy5-mRNA experiments, the 488-nm laser was adjusted to 20/50 mW and the 638-nm laser to 20/100 mW. In most cases, samples were diluted 1:50 (v/v) with PBS or aptamer detection buffer, except for the samples that were incubated with Exo-Fect^TM^ reagent or naked FAM-siRNA, which were diluted 1:80 (v/v) with PBS due to higher fluorescence background signals. Data analysis was performed using the NanoFCM Professional Suite software (version 1.8, NanoFCM Co., Ltd.), with manual threshold adjustments for each experiment to ensure comparability between tested conditions and measurement days. Threshold settings are detailed in table S3. Subsequent data analysis was conducted using FlowJo^TM^ software (version 10.4.2, BD Biosciences, USA).

### Single-particle immunoaffinity pull-down assays

Single-particle level quantification of the surface-protein expression profiles of EVs and HEVs as well as determination of FAM-siRNA colocalization in these particles were performed on an optofluidic platform based on a negative pressure-driven microfluidic biosensing chip, and imaged on a custom-built time-multiplexed single molecule fluorescence and interferometric scattering microscope (fig. S10 and Supplementary Materials). Briefly, NPs expressing the targeted surface proteins were immobilized to the surface *via* an antibody-functionalized supported lipid bilayer immunoaffinity assay prepared directly within the microfluidic chip. HEVs, LCNPs and EVs at a concentration of 1-2 × 10^10^ particles/mL were introduced to the immunoassay functionalized chip at a flow rate of 5 mL/min for 10 min. After rinsing unbound particles with buffer, the sensor was imaged sequentially in fluorescent and label-free channels across an area of 0.1 mm^2^. Single particles from each imaging channel were then colocalized from which the total amount of immobilized particles and corresponding fluorescent fraction were determined (fig. S11 and Supplementary Materials).

### CD73 and CD26 enzymatic activity

The enzymatic activities of two EV membrane proteins, CD73 and CD26, were evaluated following HEV formation and compared to EVs that underwent transfection with the Exo-Fect^TM^ reagent or were treated with enzyme inhibitors (40 µM APCP for CD73 or 1 µM linagliptin for CD26). HEV formation was carried out at both 4 and 37 °C for 30 min in HEBS (pH 7.4) at an EV:LCNP particle number ratio of 1:1.

For CD73 activity measurements, an absorbance-based malachite green assay was employed, as previously described (*38, 52*). Briefly, 0.1 μg of EV proteins as determined by the Micro BCA^TM^ Kit, were suspended in 60 µL of 10 mM Tris buffer (pH 7.4) and transferred in a 96-well plate. Subsequently, 30 µL of an 800 µM AMP solution in 10 mM Tris buffer was added to the wells, and the mixtures were incubated for 10 min at 25 °C. The enzymatic reaction was terminated by adding 40 µL of a color reagent (0.034% (w/v) malachite green, 1.55% (w/v) ammonium molybdate tetrahydrate, and 0.0625% (v/v) Tween^®^ 20). The color reaction was allowed to develop for 1 h at 25 °C, and absorbance was measured at 620 nm using the Tecan Infinite^®^ M200 plate reader (Tecan). Background-corrected absorbance values were normalized with values obtained from EVs subjected to identical hybridization conditions.

CD26/DPP4 enzymatic activity was assessed using the commercially available DPPIV-Glo^TM^ Protease Assay Kit (Promega), following manufacturer’s instructions. In brief, 0.08 μg of EV proteins in 25 µL of HEBS were loaded into white 384-well plates and mixed 1:1 (v/v) with DPPIV-Glo^TM^ reagent. Luminescence was measured after 30 min of incubation at 25 °C using the Tecan Infinite^®^ M200 Pro plate reader (Tecan). Background-corrected values were normalized to the luminescence signals obtained from EVs subjected to identical hybridization conditions.

### Nuclease protection assays

To investigate the protection of exogenous NA cargo against nuclease degradation, 0.5 µg of naked Broccoli RNA aptamer or loaded in either LCNPs or HEVs were mixed with 5 µg/mL of RNase A in Broccoli aptamer buffer to reach a total volume of 27 µL. After the addition of 3 µL of 10x Broccoli aptamer detection buffer (20 mM HEPES, 150 mM KCl, 1 mM MgCl_2_, and 200 µM DFHBI-1T), fluorescence signals were recorded at 37 °C within a sealed, black 384-well plate using the Tecan Spark^®^ microplate reader. Measurements were taken every hour for a total of 24 h, with excitation and emission settings at 472 ± 5 nm and 507 ± 5 nm, respectively.

To assess the protection of endogenous NA cargo within EVs, approximately 7 × 10^11^ EVs alone or hybridized with empty LCNPs (1:1 particle number ratio; at 37 °C for 30 min) were treated with 5 µg/mL RNase A at 37 °C for 1 h. RNA was then isolated using the mirVana^TM^ miRNA Isolation Kit (Thermo Fisher Scientific), following manufacturer’s instructions, and eluted with 50 μL of ultrapure water. The samples were subsequently snap-frozen, lyophilized (Alpha 2-4 LSC, Christ, Germany), and redissolved in 2 μL of ultrapure water. The quality of the extracted RNA was assessed *via* automated electrophoresis using the RNA ScreenTape assay on an Agilent 2200 TapeStation system and analyzed with the TapeStation Analysis software (all Agilent Technologies, USA).

### Uptake and endosomal escape

Uptake studies were conducted in both HeLa cells and HeLa cells stably expressing the galectin-8 mRuby3 fusion protein (HeLa-Gal8-mRuby3).

For flow cytometry experiments, 15,000 HeLa cells were seeded in 48-well plates. After overnight incubation at 37 °C and 5% CO_2_, cells were incubated with HEVs/LCNPs dual-labeled with Atto 488 DOPE and Cy5-siRNA. The final siRNA concentration per well was 20 nM (∼4 × 10^11^ particles/mL). After 4 and 24 h of incubation, medium was removed, and cells were washed twice with FBS to remove unspecific binding. Cells were washed with PBS and subsequently trypsinized and resuspended in full DMEM. After centrifugation at 300 × *g* for 5 min, cells were washed once more with PBS and stained with the Zombie Violet^TM^ Fixable Viability Kit (Biolegend, USA) for 15 min following manufacturer’s instructions. The cells were pelleted at 300 × *g* for 10 min and resuspended in 100 μL of ice-cold flow cytometry buffer (PBS with 0.5% (w/v) BSA and 2 mM EDTA). The MFI of 10,000 cells was measured with CytoFLEX S flow cytometry instrument (Beckman Coulter). The gain settings were set to 50 for the FITC channel (blue 488-nm laser), 50 for the PB450 channel (violet 405-nm laser) and 250 for the APC channel (red 638-nm laser). The FlowJo^TM^ software was used for gating and data analysis.

For confocal microscopy, 70,000 HeLa cells, or HeLa-Gal8-mRuby3 cells were seeded in 24-well plates on sterile 12 mm coverslips. Following an overnight incubation at 37 °C and 5% CO_2_, cells were incubated with Cy5-siRNA-loaded HEVs or LCNPs for 4 h with a final siRNA concentration of 100 nM per well (∼1.6 × 10^11^ particles/mL). Free Cy5-siRNA (equivalent concentration) and PBS were employed as negative controls. CellTrace^TM^ Far Red-labeled EVs were added at a concentration of 20 μg of EV proteins per well (∼3.4 × 10^10^ particles/mL). Subsequently, cells were washed thrice with PBS and fixed with a 2% (w/v) methanol-free PFA solution in PBS for 30 min at 25 °C. To visualize cell nuclei, Hoechst 33342 stain was applied for 8 min at 25 °C, while actin filaments were stained with Phalloidin iFluor488 (Abcam, UK) for 90 min at 25 °C. Finally, coverslips were mounted using Mowiol^®^ 4-88 (Sigma-Aldrich) and imaged on a Leica SP8 confocal microscope with a HC PL APO 63x/1.40 OIL CS2 objective (all Leica Microsystems, Germany). Images were processed using Fiji software.

### GFP knockdown experiments

For siRNA transfection experiments, 30,000 HeLa-GFP cells per well were seeded in a 24-well plate. Following a 24 h incubation at 37 °C and 5% CO_2_, cells were treated with LCNPs or HEVs containing siGFP or siCTRL. Additionally, a positive control was prepared using Lipofectamine^TM^ RNAiMAX (Thermo Fisher Scientific) and siGFP according to manufacturer’s instructions. Wells treated with PBS and naked siGFP served as negative controls. Samples were prepared in 100 µL of Opti-MEM^TM^ and added to 400 µL of 10% (v/v) FBS-containing DMEM, resulting in a final siRNA concentration of 10 nM per well (∼2.5 × 10^10^ particles/mL) and an FBS concentration of 8% (v/v). After 48 h of incubation, GFP expression was qualitatively assessed using fluorescence microscopy with a Leica DMI6000 B epifluorescence microscope (Leica Microsystems) at 10x magnification, gain settings set to 2, and an exposure time of 1 ms. Subsequently, cells were washed twice with PBS, trypsinized and resuspended in full DMEM. Cells were then centrifuged at 300 × *g* for 5 min, and cell pellets were washed with PBS before being treated with the Zombie NIR^TM^ Fixable Viability Kit (Biolegend) following the manufacturer’s instructions. After 15 min incubation, cells were pelleted at 300 × *g* for 10 min and resuspended in 120 µL ice-cold flow cytometry buffer for analysis. The GFP expression of 10,000 cells was analyzed using a CytoFLEX S flow cytometry instrument (Beckman Coulter) with gain settings adjusted to 2 for the FITC channel (blue 488-nm laser) and 10 for the APC-A750 channel (red 638-nm laser). The MFI was normalized (in %) to values obtained from untreated HeLa-GFP cells and the data analysis was performed with the FlowJo^TM^ software. Additional experiments were performed under challenging cell culture conditions, increasing the FBS concentration to 80% (v/v) FBS and reducing the contact time with cells to 4 h. GFP expression was analyzed 48 h after initial sample addition using flow cytometry as outlined above.

### Luciferase expression experiments

For mRNA transfection experiments, 5,000 HeLa cells per well were seeded in 96-well plates. After 24 h of incubation at 37 °C and 5% CO_2_, the culture medium was replaced with either 90 µL of DMEM containing 10% (v/v) FBS or pure FBS. Subsequently, 10 to 12 μL (depending on the EV batch) of FLuc-mRNA-loaded LCNPs and HEVs in PBS were added. The actual mRNA concentration in the LCNP formulations was determined using the Quant-it^TM^ RiboGreen RNA Assay Kit before each experiment. The final FBS concentration in the wells was approximately 9 or 90% (v/v), with mRNA doses ranging from 0.1 to 100 ng/well (corresponding to concentrations of ∼0.3 pM to 0.3 nM and ∼1.8 × 10^8^ to 1.8 × 10^11^ particles/mL). A positive control was prepared using Lipofectamine^TM^ MessengerMAX^TM^ (Thermo Fisher Scientific) with FLuc-mRNA, following the manufacturer’s instructions, with the only difference being that particle formation was carried out in PBS instead of OptiMEM^TM^. PBS and naked FLuc-mRNA served as negative controls. After 24 h of incubation, the cells were washed twice with DPBS, and luciferase expression was assessed using the Pierce^TM^ Firefly Luciferase Glow Assay Kit (Thermo Fisher Scientific). In brief, cell lysates were prepared with 40 µL of 1x cell lysis buffer, and 5 µL of the lysates were transferred into a white 384-well plate. Then, 25 µL of the working reagent (firefly glow assay buffer with 1x D-luciferin and 1x Pierce^TM^ firefly signal enhancer) was added, and luminescence was measured after 10 min of incubation at 25 °C using the Tecan Spark^®^ microplate reader. Luminescence readings were normalized to the amount of loaded protein, as determined with the Micro BCA^TM^ Protein Assay Kit, and the background signal of untreated cells was subtracted from the treatment conditions.

### Statistical analysis

All experimental findings in this study are from at least 3 independent experiments and data is presented as mean +/- standard deviation (s.d.), as indicated in each figure legend. Statistical significance between two groups was calculated with an unpaired Student’s t-test. For mRNA transfection experiments, the comparison between HEV and LCNP conditions was conducted with an unpaired Student’s t-test, excluding the positive and negative controls from the statistical analysis. Statistically significant differences between multiple groups were calculated by one-way or two-way ANOVA followed by Tukey’s post-hoc test. Statistical analyses and curve fitting was performed using GraphPad Prism software (version 8.2.0, GraphPad Software Inc., USA). A p value of < 0.05 was considered statistically significant.

## Acknowledgments

We would like to thank Stephan Handschin and Miroslav Peterek (Scientific Center for Optical and Electron Microscopy (ScopeM), ETH Zurich) for performing EM measurements. We would also like to thank Dr. Claudia Dumrese and Dr. Philipp Schätzle (Cytometry Facility, University of Zurich) for their help with NanoFCM measurements. A special thanks to Dr. Felix Gloge and Wyatt Technology for providing access to the DynaPro Plate Reader. Dr. Freideriki Michailidou, Dr. Angela Steinauer and Prof. Donald Hilvert are acknowledged for sharing Broccoli RNA aptamer production/purification protocols and for access to their laboratory equipment. Dr. Anna Knörlein is acknowledged for analyzing RNA aptamers with LC-MS. Further, we would like to thank David Gremmelspacher and Prof. Nicola Aceto for their help with TapeStation measurements and access to the instrument. Prof. Klaus Eyer and Dr. Ines Lüchtefeld are acknowledged for the use and their help with the syringe pump system. Finally, we would like to thank Jasmin Maier and Fanxi Meng for their experimental support, as well as Dr. Britta Hettich for her input on figure illustrations.

## Author contributions

Conceptualization: JB, EM, JCL

Methodology: JB, PR, VM, JOA

Investigation: JB, PR, VM, SG, FB, BMQ, JOA, EM

Visualization: JB, PR, VM, JOA

Supervision: JOA, JCL

Writing—original draft: JB

Writing—review & editing: JB, PR, VM, SG, FB, BMQ, JOA, EM, JCL

## Data and materials availability

All data are available in the main text or the supplementary materials.

