## Supplementary Materials for "Loading of extracellular vesicles with nucleic acids via hybridization with sponge-like lipid nanoparticles"

Johannes Bader *et al.*

**This PDF file includes:**

Supplementary Methods
Figs. S1 to S15
Tables S1 to S3

**Other Supplementary Materials for this manuscript include the following:**

Movies S1 and S2

**Supplementary Methods**

Liquid chromatography mass spectrometry (LC-MS)

LC-MS for Broccoli aptamer samples was performed on an Acquity OST C18 column (Waters, USA) using an Agilent 1200/6130 system (Agilent Technologies, USA). The aptamer samples were separated by a 0.4 M hexafluoroisopropanol (Fluorochem, UK) and 15 mM trimethylamine (Sigma-Aldrich, USA) in water (solvent A) and methanol (Thermo Fisher Scientific, USA) (solvent B) gradient (5-80 % B in 10 min) at a flow rate of 0.3 mL/min.

Molecular fingerprinting reagents

Bovine serum albumin (BSA, Sigma-Aldrich, USA) solutions were prepared in phosphate buffered saline (PBS, pH 7.4, Sigma-Aldrich). Neutravidin (Thermo Fisher Scientific, USA) was diluted to a concentration of 0.02 mg/mL (0.3 μM) in PBS with 1% (w/v) BSA. Biotinylated antibodies anti-CD63, and anti-CD9, from Ancell (USA); and anti-CD26, anti-CD73 and Mouse IgG1 κ-isotype control from Biolegend (USA) were prepared to a concentration of 0.05 mg/mL (0.33 μM) in PBS with 1% (w/v) BSA for all experiments. A custom AH peptide with the following sequence: SGSWLRDVWDWICTVLTDFKTWLQSKLDYKD was synthesised by Proteogenix (France). A stock solution of 1 mg/mL was prepared by dissolving the lyophilised peptide in Milli-Q water according to manufacturer's recommendation. This stock solution was aliquoted and stored at -20 °C for up to one month. For all experiments, the peptide stock solution was diluted to 0.45 mg/mL (200 μM).

Liposome preparation for microfluidic *in situ* supported lipid bilayer formation

Liposomes were prepared using the extrusion method with 99:1 (mol%) lipid composition of 1-palmitoyl-2-oleoyl-*sn*-glycero-3-phosphocholine (16:0-18:1, POPC) (Avanti Polar Lipids, USA) to 1,2-dioleoyl-*sn*-glycero-3-phosphoethanolamine-*N*-(cap biotinyl) (18:1 biotin-DOPE) (Avanti Polar Lipids) by first mixing the two lipid stock solutions in chloroform. The mixed lipid solution was subsequently dried under a nitrogen stream and then placed under vacuum for 24 h. The dried lipid film was then rehydrated to a concentration of 5 mg/mL in Tris buffer (100 mM, pH 7.4, Sigma-Aldrich) and vortexed for 2 min. The hydrated lipid stock solution was then extruded sequentially through polycarbonate membranes of 100, 50 and 30 nm using Avanti Polar Lipids Mini-Extruder. For each sequential extrusion, the lipid solution was passed 21 times through the polycarbonate membrane. For all experiments, the liposome solution was prepared at a concentration of 1 mg/mL, and used within 3 days of preparation. Finally, 5 μL of a 500 mM $CaCl_2$ (Millipore, USA) solution in Milli-Q water was added to 500 μL of the 1 mg/ml liposome dilution.

Fabrication of the microfluidic chips for molecular fingerprinting

The microfluidic chips used for the immunoaffinity pulldown assays were fabricated using standard soft lithography processes. A microfluidic chip design based on negative pressure was used. The sample was injected by simple pipetting to a chip inlet area and flow was adjusted by applying vacuum at the outlet. The outlet of the sensing area chip was connected to a resistance chip to adjust the flow rates to ideal pressure conditions in the range of -100 to -700 mbar. Microfluidic chip moulds were made on silicon wafers using a laser writer (at 365 nm, μMLA, Heidelberg Instruments, Germany) with SU8 1060 photoresist (Gersteltec, Switzerland) resulting in channel thicknesses of approximately 100 μm for both moulds. The microfluidic chips are made from polydimethylsiloxane (PDMS, SYLGARD™ 184 Silicone Elastomer Kit, Dow Europe, Germany) mixed at a ratio of 10:1 (w/w) polymer to curing agent. The PDMS was drop casted onto the wafers and then degassed under vacuum until all bubbles were removed and baked in a convection oven at 80 °C for 1 h. The cured PDMS was peeled off the mould and holes for sample injection and pinning were punched. The PDMS chips and cleaned glass coverslips (24 × 40 $mm^2$, 0.17 mm, Karl Hecht, Germany, for the sensing chip; SuperFrost® microscope slides, VWR, USA, for the resistance chip) were treated with oxygen plasma (13.56 MHz, 10.5 L, Atto, Diener electronic, Germany) for 1 min (300 W, 1.5 sccm $O_2$ flow rate) before binding together and baking for 1 h in an 80 °C oven.

Molecular fingerprinting immunoassay

The chip was primed and washed with 150 µL PBS at 30 µL/min. Then, 20 µL of the 1 mg/mL liposome stock solution was injected into the channels until a bilayer had visibly formed. Once bilayer formation had occurred, the channels were rinsed with 150 µL MilliQ-water for 5 min to remove any excess unbound liposomes. After rinsing, 20 µl of the AH peptide solution was introduced into the channel. The bilayer was then rinsed with 150 µl PBS for 5 min. This results in a fully formed bilayer coating. Next, 20 µL of the 0.02 mg/mL Neutravidin solution was flowed into the channel and incubated for 30 min. The channel was again rinsed with 150 µL PBS for 5 min and 20 µL of the chosen capture antibody solution was flowed into the channels and incubated for 30 min. A final rinsing step with 150 µL PBS for another 5 min completed the passivated and functionalized surface ready for immunoaffinity based capturing. Reference scans were performed upon completion of the surface functionalization of each channel. For the fingerprinting assay, LCNP, EV and HEV samples were diluted to a target concentration in the range of 1-2 × $10^{10}$ particles/mL on the day of experiment and were introduced at a flow rate of 5 µL/min for 10 min. Subsequently, the channels were washed for 5 min with PBS at a flow rate of 30 µL/min before performing imaging scans, hereby termed sample scans, over the same area as those acquired during the reference scans.

Correlative fluorescence and interferometric scattering microscope setup

The custom-built optical system was based on a common-path digital holographic microscope operating in reflection, whereby illumination and imaging arms were separated by a single 50:50 beamsplitter plate (BSW27, Thorlabs, USA) and all optics were arranged in a 4f configuration. The illumination arm was composed of two different excitation sources corresponding to the label-free and fluorescence channels, respectively. For off-resonant illumination, *i.e.* label-free channel excitation, a 617 nm light emitting diode (M617F2 LED, Thorlabs) was coupled into a 200 µm multi-mode fibre (M25L02, Thorlabs) and subsequently outcoupled with a 6.24 mm aspheric lens (A110TM-A, Thorlabs). For fluorescence excitation, a 488 nm diode laser (Lasertack, Germany) was coupled into a 150 µm × 150 µm square multimode fibre (M103L04, Thorlabs) and outcoupled with a 20x, 0.40 NA air objective (RMS20X, Thorlabs). Light outcoupled from both fibres were combined along the same illumination path with a long-pass dichroic mirror (DMLP567, Thorlabs) and relay imaged onto the sample plane formed by a 1.46 NA oil immersion objective (APON 60XOTRIF, Olympus, Japan) via a 1:1 imaging system, composed of two 300 mm achromatic doublet lenses (AC508-300A, Thorlabs). Under this optical arrangement, the label-free channel was illuminated with a flat-top profile of 89.5 µm diameter and $NA_{illumination} = 0.5$; whereas the fluorescence channel was illuminated with a square flat-top profile 50 µm in length.

For the imaging arm, light collected from the sample by the objective and reflected off the 50:50 beamsplitter was first filtered by a multibandpass fluorescence filter (87-244, Edmund Optics, UK) and subsequently imaged onto a scientific CMOS camera (C11440-22CU, 6.5 µm pixels,

Hamamatsu, Japan) using a 300 mm achromatic doublet (AC508-300A, Thorlabs) resulting in a 100x magnification. The sample was mounted on a motorized XY microstage (Mad City Labs, USA) equipped with linear encoders, as well as a XYZ nanostage (Nano-LP200, Mad City Labs). The sample focus position was stabilised to within 10 nm using the backreflection from 670 nm misaligned confocal beam with a low numerical aperture of illumination (CPS670F, Thorlabs).

Correlative fluorescence and label-free imaging

For all optofluidic-based experiments, we measured a power at the sample between 0.45-0.50 mW and 4.8-5.0 mW equivalent to an irradiance of 0.07-0.08 $\mu W/\mu m^2$ and 1.92-2.00 $\mu W/\mu m^2$ for the label-free and fluorescence channels, respectively. During acquisition, a field of view of 66.6 μm × 66.6 μm corresponding to an area of 1024 × 1024 camera pixels was recorded with an exposure time of 20 ms and a fixed frame rate of 50 Hz. To minimise data load and increase the signal-to-noise (SNR) ratio, the data were saved in the form of 20 time-averaged frames, leading to an effective time resolution of 2.5 Hz. Prior to each data acquisition, an experimental flat-field image was generated and saved. The flat-field image was produced by first collecting a stack of at least 60 time-averaged frames taken at different sample locations and same focus position, and subsequently taking the median value on a pixel-by-pixel value. This flat-field image contained inhomogeneities along the optical system and imperfections in the sample illumination.

Image processing pipeline for the molecular fingerprinting assay

Label-free images were first normalised to the average camera counts in the background. Next, the normalised label-free images were flat-field corrected, by division, to remove inhomogeneities attributed to the optical system and sample illumination. Diffraction-limited spots were segmented from the image based on their local SNR ratio. Namely, only diffraction-limited spots satisfying the following two selection criteria were considered, a) pixel-based: positive for all pixels exceeding a SNR threshold of 3.5; and b) clustering-based: positive if there were a minimum of 4 pixels exceeding the SNR threshold within a 3 × 3 $pixel^2$ area. The resulting positively identified diffraction-limited spots were subsequently localised with sub-pixel precision using the radial symmetry centres algorithm where the lateral position, signal contrast and the integrated signal contrast, were stored for further processing. Total particle counts captured during the immunoaffinity assay were determined by taking the difference between the sample and reference scans. Finally total particle counts for all markers were normalised to the counts of CD73+ samples.

To correlate the fluorescence signal to the label-free one from an identified particle, the fluorescence images were first flat-field corrected, by subtraction, to remove background counts and inhomogeneities in the illumination. Next, the region of interests corresponding to particle signatures in the label-free channel were colocalized in the fluorescence image. The region of interest was to 9 × 9 $pixel^2$ from which the total fluorescent counts were determined. Only particles with fluorescent counts at least 3 standard deviations greater than the background fluorescence were assigned as positively fluorescently labelled (fluorescent fraction). The loading efficiency was then computed as the ratio of the fluorescent fraction to the total number of particles identified in the label-free channel.

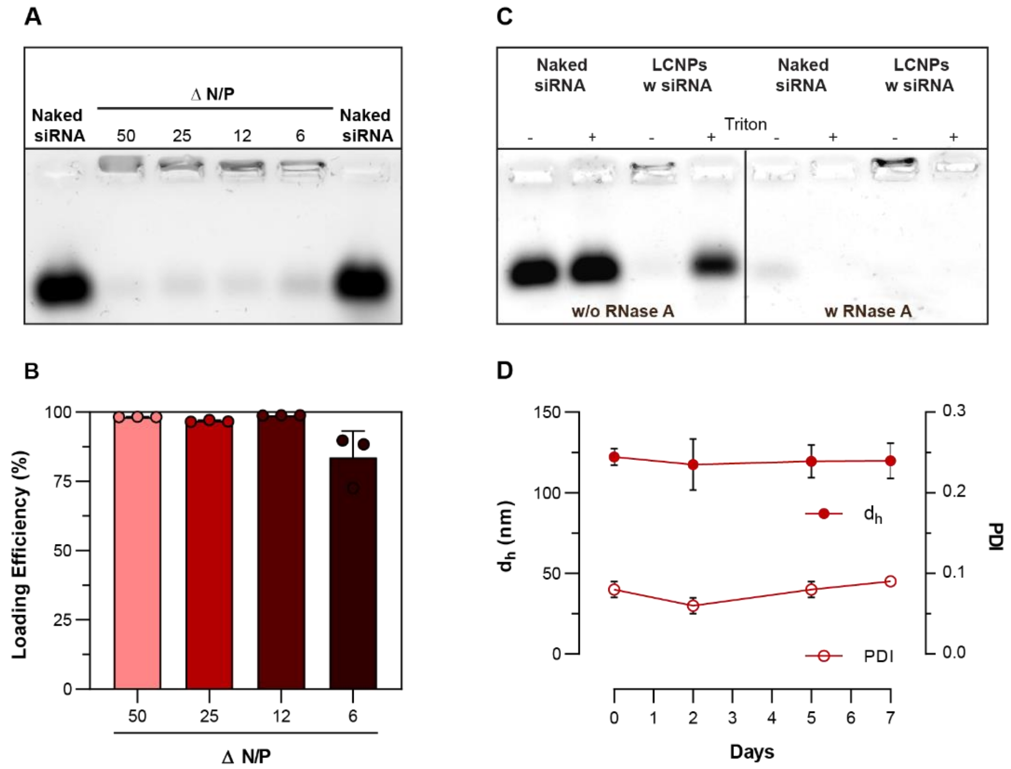


**fig. S1. Loading of siRNA in LCNPs and protection against RNase degradation.**

(**A**) Representative EMSA gel image of LCNP formulations with varying N/P ratios. (**B**) Loading efficiency of LCNP formulations with varying NP ratios as determined with the RiboGreen assay (n = 3) (data derives from three independent samples with single measurements). (**C**) Representative EMSA gel image of naked siRNA or siRNA loaded in LCNPs (N/P 25) in the presence of Triton™ X-100 and/or RNase A (5 µg/mL). (**D**) $d_h$ and PDI of siRNA-loaded LCNPs (N/P 25) over a week of incubation at 4 °C (n = 3). Data is expressed as mean + s.d. of n independent experiments.

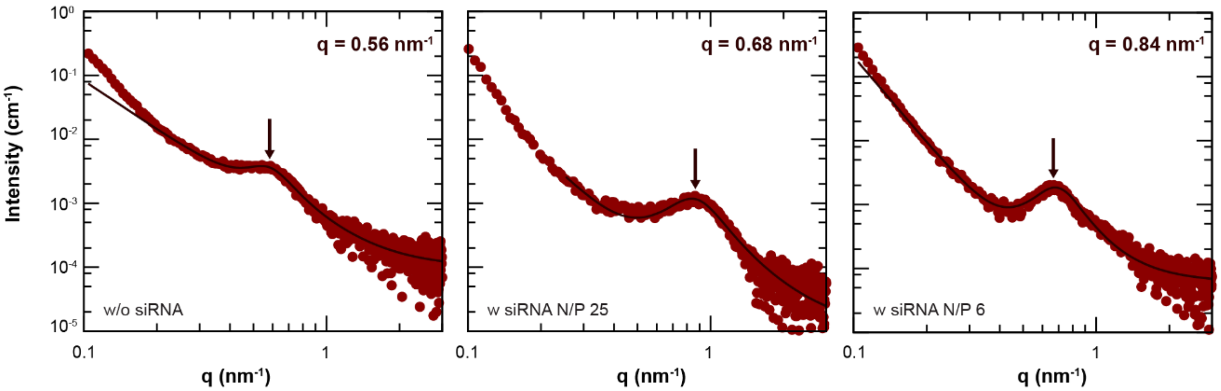


**fig. S2. Lorentzian model fitting of LCNP SAXS curves.**

Peak positions of the first structure peaks of LCNPs at pH 7.4 with (N/P 25 and 6) or without siRNA were determined with a Lorentzian broad peak model function in a q-range between 0.25 and 4 $nm^{-1}$.

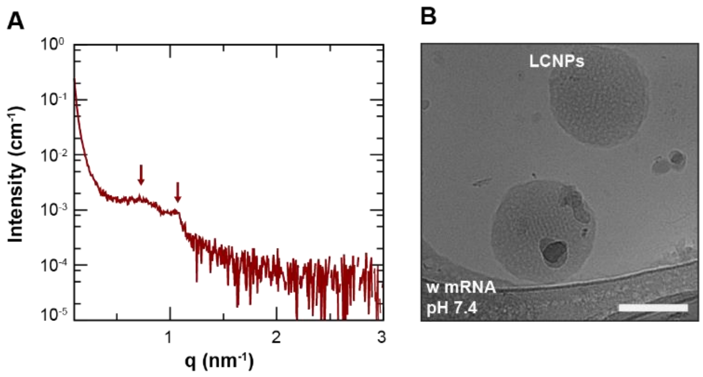


C

| | LCNPs (w mRNA) |
|---|---|
| $d_h$ (nm) | 129 ± 16 |
| PDI | 0.11 ± 0.03 |
| ζ-potential (mV) | 17.1 ± 4.5 |
| LE (%) | 92.3 ± 0.5 |

**fig. S3. Physicochemical characterization of mRNA-loaded LCNPs.**

(**A**) SAXS profile of mRNA-loaded LCNPs (N/P 25) at pH 7.4. (**B**) Representative cryo-TEM image of mRNA-loaded LCNPs (N/P 25) at pH 7.4; scale bar 100 nm. (**C**) Summary of physicochemical properties of mRNA-loaded LCNPs (N/P 25) (n = 3); LE = loading efficiency (LE data derives from three independent samples with single measurements). Data is expressed as mean ± s.d. of n independent experiments.

### S3. Supplementary discussion on SAXS analysis of mRNA-loaded LCNPs:

A broad peak at q = 0.75 $nm^{-1}$ (↓) is visible, suggesting that similar to siRNA-containing LCNPs (Fig. 2B) non-lamellar sponge ($L_3$) phases formed in the presence of mRNA (fig. S3A). This broad peak overlaps with a sharper one at q = 1.05 $nm^{-1}$ (↓), indicating the formation of domains with a narrower inter-lipid bilayer spacing, potentially due to the presence of mRNA in the aqueous compartments.

A

| | DOPC-NPs (w/o siRNA) | | DOPC-NPs (w siRNA) | |
|---|---|---|---|---|
| $d_h$ (nm) | 117 ± 11<br>105 ± 10 | pH 5<br>pH 7.4 | 123 ± 27<br>101 ± 5 | pH 5<br>pH 7.4 |
| PDI | 0.10 ± 0.01<br>0.16 ± 0.04 | pH 5<br>pH 7.4 | 0.09 ± 0.03<br>0.15 ± 0.05 | pH 5<br>pH 7.4 |
| ζ-potential (mV) | 46 ± 4.4<br>21 ± 1.3 | pH 5<br>pH 7.4 | 36 ± 10.4<br>22 ± 0.04 | pH 5<br>pH 7.4 |
| LE (%) | | | 97.0 ± 0.89 | |

| | DOTMA-NPs (w/o siRNA) | | DOTMA-NPs (w siRNA) | |
|---|---|---|---|---|
| $d_h$ (nm) | 55 ± 7<br>66 ± 8 | pH 5<br>pH 7.4 | 69 ± 14<br>70 ± 12 | pH 5<br>pH 7.4 |
| PDI | 0.09 ± 0.02<br>0.09 ± 0.01 | pH 5<br>pH 7.4 | 0.13 ± 0.03<br>0.13 ± 0.03 | pH 5<br>pH 7.4 |
| ζ-potential (mV) | 54 ± 1.4<br>41 ± 10.5 | pH 5<br>pH 7.4 | 49 ± 5.4<br>48 ± 0.5 | pH 5<br>pH 7.4 |
| LE (%) | | | 100.1 ± 0.23 | |

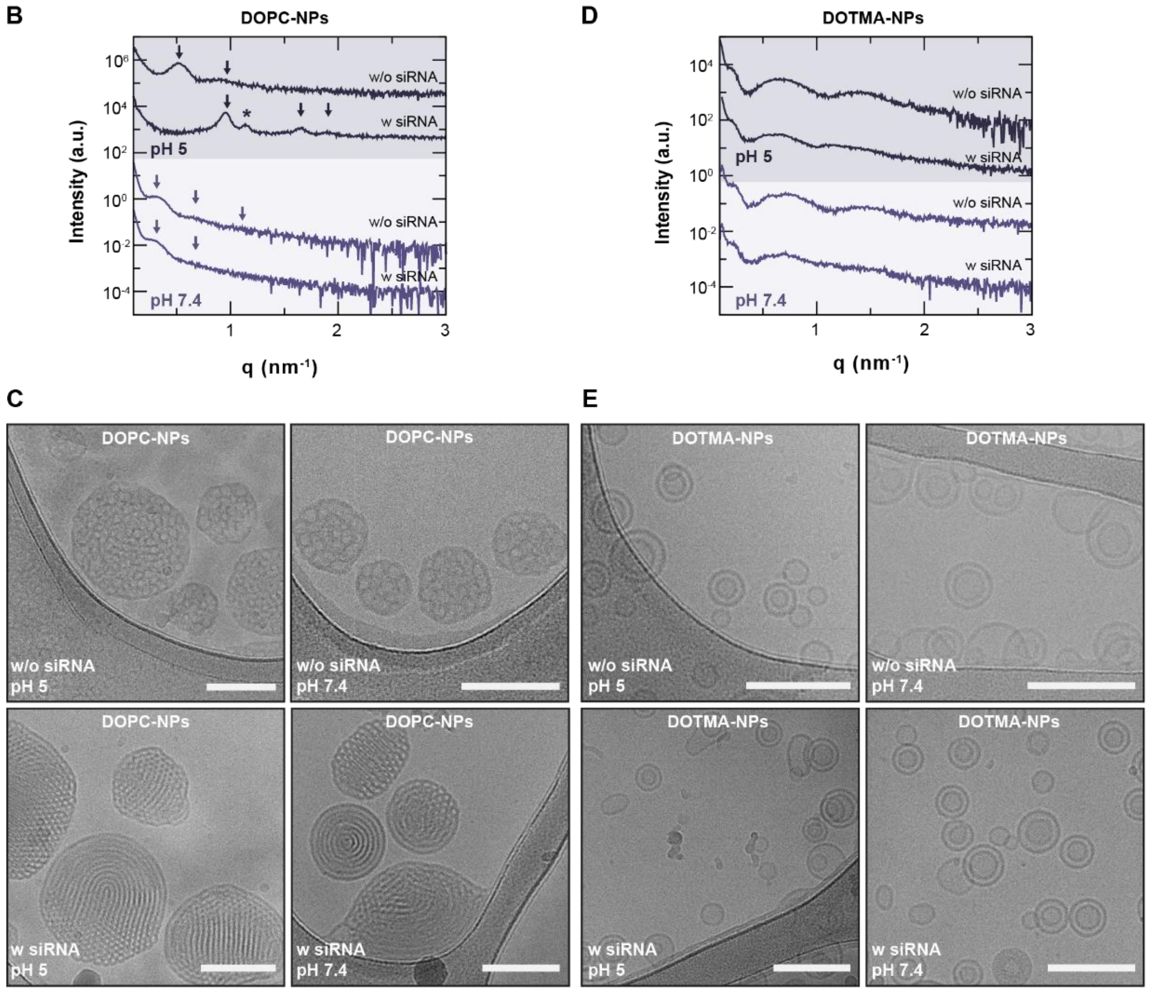


**fig. S4. Physicochemical characterization of DOTMA- and DOPC-NPs.**

Structural properties of LCNPs can be altered by substituting the inverse cone-shaped lipids DOPE or MC3 (18 or 60 mol%) with cylindrical lipids DOPC or DOTMA (18 or 60 mol%). (**A**) Summary of the physicochemical properties of unloaded and siRNA-loaded DOTMA- and DOPC-NPs (N/P 25) at pH 5 and 7.4 (n = 3); LE = loading efficiency. Data is expressed as mean ± s.d. of n independent experiments. (**B**) SAXS scans of DOPC-NPs at two different buffer pH values (5 and 7.4) with (N/P 25) or without siRNA. (**C**) Representative cryo-TEM images of DOPC-NPs at pH 5 and 7.4 with (N/P 25) and without siRNA; scale bars 100 nm. (**D**) SAXS scans of DOTMA-NPs at two different buffer pH values (5 and 7.4) with (N/P 25) or without siRNA. (**E**) Representative cryo-TEM images of DOTMA-NPs at pH 5 and 7.4 with (N/P 25) and without siRNA; scale bars 100 nm.

**S4. Supplementary discussion on SAXS analysis of DOPC- and DOTMA-NPs:**

At pH 7.4, SAXS scans of unloaded DOPC-NPs revealed a broad peak maximum at a q value of 0.33 $nm^{-1}$ (↓), indicating a structure with a repeat distance of approximately 19 nm (fig. S4B). Additionally, two weaker oscillations at q values of approximately 2 to 4 times of the main structure peak were detected (↓). The repeat distance of 19 nm was much larger than the expected thickness of a bilayer or compared to $L_3$ phases and suggests that there is a higher water content within the system. Taken together the SAXS profile aligns more with that of multivesicular liposomes, suggesting that the internal structure is made of small vesicular domains. Upon the addition of siRNA, the structure of DOPC-NPs appeared more disordered and only the first correlation peak could be clearly observed at a q value of 0.31 $nm^{-1}$ (↓) along a second broad oscillation at around 0.62 $nm^{-1}$ (↓) (fig. S4B). At pH 5, the main structure peak of unloaded DOPC-NPs became sharper and shifted towards a higher q value of 0.52 $nm^{-1}$ (↓), suggesting a lower spacing among the lipid bilayers (~12 nm) (fig. S4B). The emergence of a second broad peak at q = 0.90 $nm^{-1}$ (↓), positioned approximately √3 times the location of the initial peak, suggests that the particles adopted a hexagonal arrangement, albeit with higher hydration compared to $H_{II}$ and $L_3$ phases found in LCNPs. Upon addition of siRNA, 3 sharp peaks at q values of 0.95 $nm^{-1}$, 1.65 $nm^{-1}$ and 1.9 $nm^{-1}$ (↓) with a q-position ratio of 1:√3:2 indicate that a hexagonal phase formed in DOPC-NPs at pH 5 (fig. S4B). An additional peak at q = 1.14 $nm^{-1}$ appeared (*), suggesting the presence of a second population similar to LCNPs (Fig. 2B).
The formulations containing DOTMA at pH 7.4 and at pH 5 showed, in both cases, a series of broad oscillations, suggesting a low degree of multilamellarity of the lipid bilayers. At both pH values, after adding siRNA, the main oscillation shifted towards lower q values, indicating that the overall thickness is increasing due to either the bilayer becoming thicker or an additional layer being stabilized (fig. S4D).

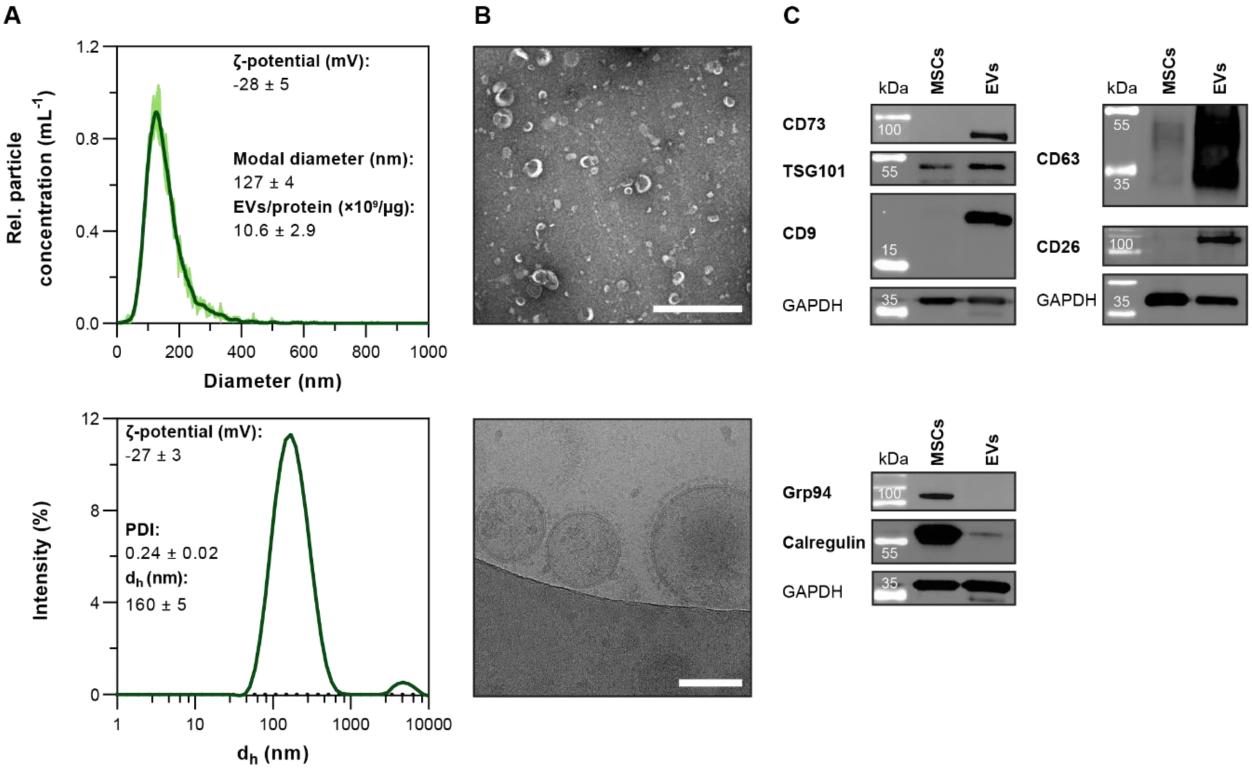


**fig. S5. Characterization of MSC-derived EVs.**

(**A**) Size distribution profiles of EVs as characterized by NTA (top) and DLS (bottom) (n = 3). Data is expressed as mean ± s.d. of n independent experiments. (**B**) Representative TEM (top) and cryo-TEM image (bottom) of EVs; scale bars 1 µm (top) and 100 nm (bottom). (**C**) Expression levels of EV marker proteins (CD9, CD63 and TSG101), MSC-specific membrane proteins (CD73 and CD26) and cellular impurity markers (calregulin and Grp94) in cell and EV lysates analyzed by western blotting. GAPDH serves as loading control. One representative blot is shown.

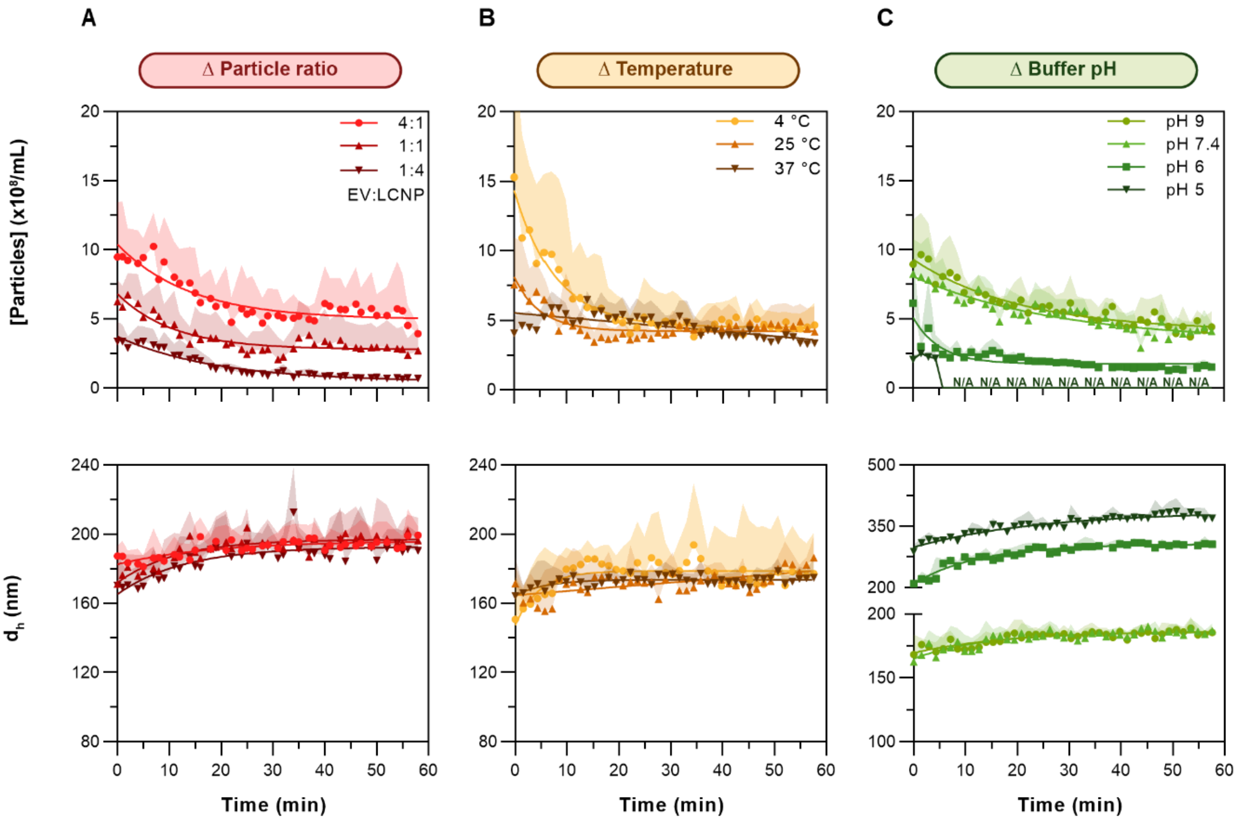


**fig. S6. DLS/SLS time-course experiments.**

Particle concentration and $d_h$ of mixtures of EVs and siRNA-loaded LCNPs (N/P 25) measured over a 60-min time course by SLS and DLS, respectively; (**A**) as function of EV:LCNP number ratio (4:1, 1:1 and 1:4); (**B**) as a function of temperature (4, 25 and 37 °C) at a 1:1 EV:LCNP particle number ratio; and (**C**) as a function of pH (9, 7.4, 6 and 5) at a 1:1 EV:LCNP particle number ratio (n = 3-5). Data is expressed as mean + s.d. of n independent experiments.

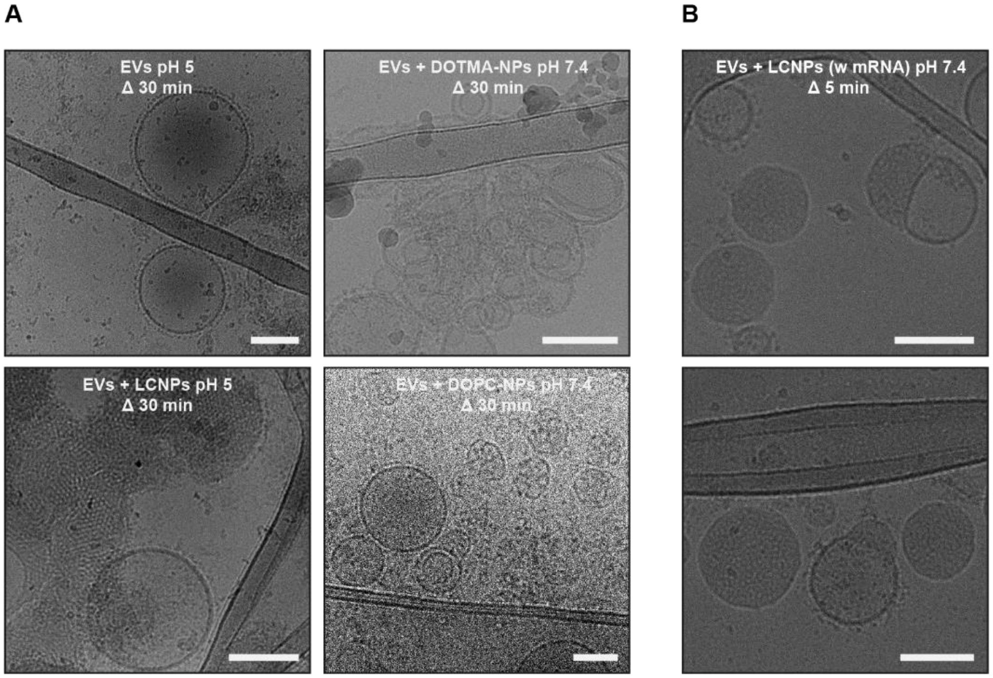


**fig. S7. Cryo-TEM imaging of mixtures of EVs with various lipid nanocarriers.**

(**A**) Representative cryo-TEM images of EVs at pH 5 and mixtures of EVs and siRNA-loaded LCNPs (N/P 25) at pH 5; EVs and siRNA-loaded DOTMA-NPs (N/P 25) at pH 7.4 and EVs and siRNA-loaded DOPC-NPs (N/P 25) at pH 7.4. Samples were mixed at particle number ratios of 1:4 and imaged after 30 min of incubation at 37 °C; scale bars 100 nm. (**B**) Representative cryo-TEM images of mixtures of EVs and mRNA-loaded LCNPs (N/P 25) at a particle number ratio of 1:4 after 5 min of incubation at 37 °C and pH 7.4; scale bars 100 nm.

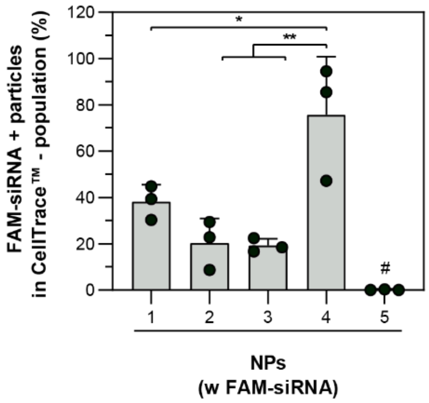


**fig. S8. Percentage of loaded fluorescent siRNA in lipid nanocarriers.**

Percentage of FAM-siRNA+ particles ae shown relative to the respective total CellTrace$^{TM}$-particle populations as determined by NanoFCM analysis (n = 3). The various FAM-siRNA-loaded NPs (N/P 25) are labeled from 1 to 5: (1) LCNPs pH 7.4; (2) LCNPs pH 5; (3) DOPC-NPs pH 7.4; (4) DOTMA-NPs pH 7.4; (5) Exo-Fect$^{TM}$ pH 7.4. Statistical significance was calculated from an ordinary one-way ANOVA with Tukey's post-hoc test; *p < 0.05 and **p < 0.01; # = *vs.* 1 *p < 0.05 and *vs.* 4 ***p < 0.001.

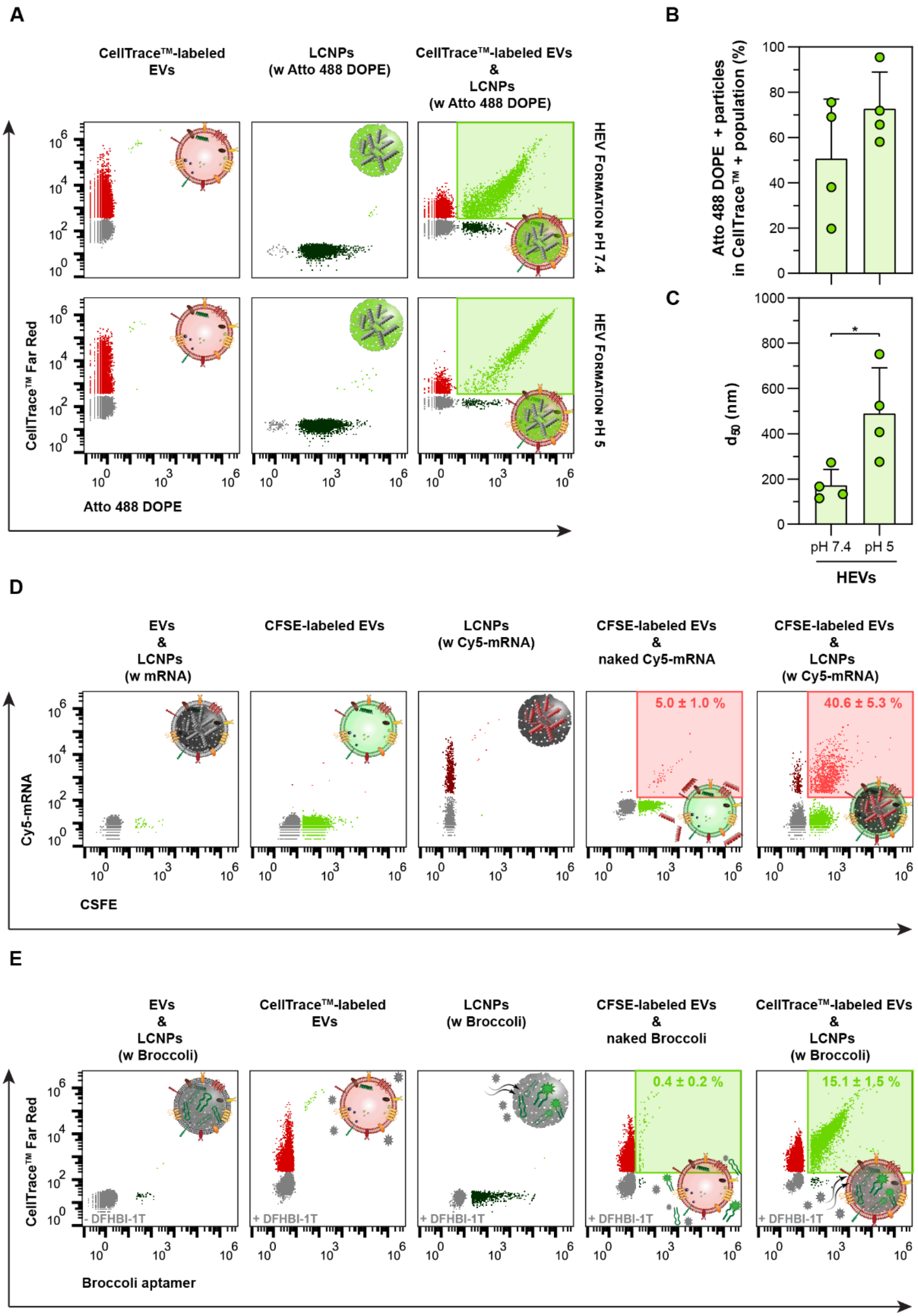
A
CellTrace™-labeled EVs
LCNPs (w Atto 488 DOPE)
CellTrace™-labeled EVs & LCNPs (w Atto 488 DOPE)
HEV FORMATION PH 7.4
HEV FORMATION PH 5
CellTrace™ Far Red
Atto 488 DOPE
B
Atto 488 DOPE + particles in CellTrace™ + population (%)
C
d50 (nm)
pH 7.4
pH 5
HEVs
D
EVs & LCNPs (w mRNA)
CFSE-labeled EVs
LCNPs (w Cy5-mRNA)
CFSE-labeled EVs & naked Cy5-mRNA
CFSE-labeled EVs & LCNPs (w Cy5-mRNA)
5.0 ± 1.0 %
40.6 ± 5.3 %
Cy5-mRNA
CSFE
E
EVs & LCNPs (w Broccoli)
CellTrace™-labeled EVs
LCNPs (w Broccoli)
CFSE-labeled EVs & naked Broccoli
CellTrace™-labeled EVs & LCNPs (w Broccoli)
0.4 ± 0.2 %
15.1 ± 1.5 %
- DFHBI-1T
+ DFHBI-1T
CellTrace™ Far Red
Broccoli aptamer

**fig. S9. Characterization of the HEV formation with fluorescent lipids and different NA cargos by NanoFCM.**

(**A**) CellTrace$^{TM}$ Far Red-labeled EVs are combined with Atto 488 DOPE-labeled (1 mol%) LCNPs (N/P 25) at particle number ratios of 1:1 and incubated at 37 °C for 30 min at pH 5 or 7.4. Representative bivariate dot plots of Atto 488 DOPE+ particles (x-axis) *vs.* CellTrace$^{TM}$+ particles (y-axis) show the emergence of a double-positive (CellTrace$^{TM}$+/Atto 488 DOPE+) particle population after coincubation of Atto 488 DOPE-labeled LCNPs with CellTrace$^{TM}$-labeled EVs as highlighted by the lime green square inset. (**B**) Percentages of the double-positive particles (CellTrace$^{TM}$+/Atto 488 DOPE+) relative to the respective total CellTrace$^{TM}$+ particle population (n = 4). (**C**) $d_{50}$ of the double-positive (CellTrace$^{TM}$+/Atto 488 DOPE+) particle population (n = 4). (**D**) CFSE-labeled EVs are combined with Cy5-mRNA-loaded LCNPs (N/P 25) at a particle number ratio of 1:1 and incubated at 37 °C for 30 min at pH 7.4. Representative bivariate dot plots of CFSE+ particles (x-axis) *vs.* Cy5-mRNA+ particles (y-axis) show the emergence of a double-positive (CFSE+/Cy5-mRNA+) particle population after coincubation of Cy5-mRNA-loaded LCNPs and CFSE-labeled EVs as highlighted by the pink square inset. Coincubation control with naked Cy5-mRNA (~0.1 µM), single-labeled and unlabeled controls are provided (n = 3). (**E**) CellTrace$^{TM}$ Far Red-labeled EVs are combined with Broccoli aptamer-loaded LCNPs (N/P 6) at particle number ratios of 1:1 and incubated at 37 °C for 30 min at pH 7.4. Representative bivariate dot plots of Broccoli aptamer+ particles (x-axis) *vs.* CellTrace$^{TM}$+ particles (y-axis) show the emergence of a double-positive (CellTrace$^{TM}$+/Broccoli aptamer+) particle population after coincubation of Broccoli aptamer-loaded LCNPs with CellTrace$^{TM}$-labeled EVs as highlighted by the lime green square inset. Coincubation control with naked Broccoli (~6 µM) and single-stained controls in the presence of the aptamer-selective dye DFHBI-1T, as well as unstained controls without DFHBI-1T are provided (n = 3). Data is expressed as mean +/± s.d. of n independent experiments. Statistical significance was calculated with a two-tailed unpaired Student's t-test; $*p < 0.05$.

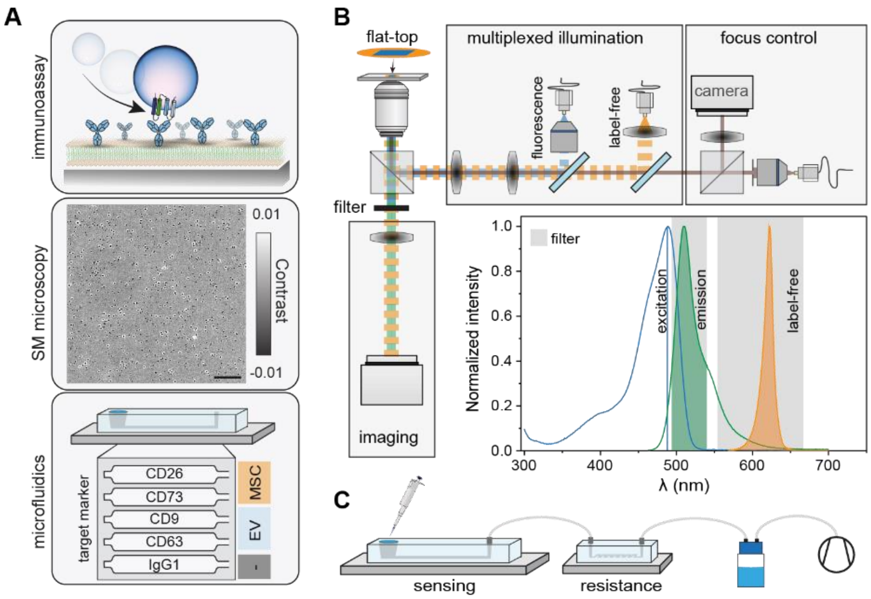


**fig. S10. Optofluidic platform for molecular fingerprinting.**

(**A**) Conceptual schematic of the optofluidic platform that combines immunoaffinity pulldown assays with single molecule (SM) microscopy, and microfluidic-based multiplexed target screening. Scale bar: 10 µm. (**B**) Schematic of the custom-built microscope setup for correlative single molecule fluorescence and interferometric scattering microscopy. Inset: fluorescence absorption and emission spectra of FAM-siRNA overlaid with the excitation light sources for fluorescent and label-free channels. Shaded regions indicate the transmission bands of the multi-bandpass filter. (**C**) Schematic of the negative-pressure driven microfluidic system architecture comprising a sensing chip, a resistance chip, and a vacuum pump. Only the sensing chip is placed on the optical microscope and reagents are introduced with a pipette and flown through the chip by applying a negative pressure; whereas the resistance chip controls the flow profile.

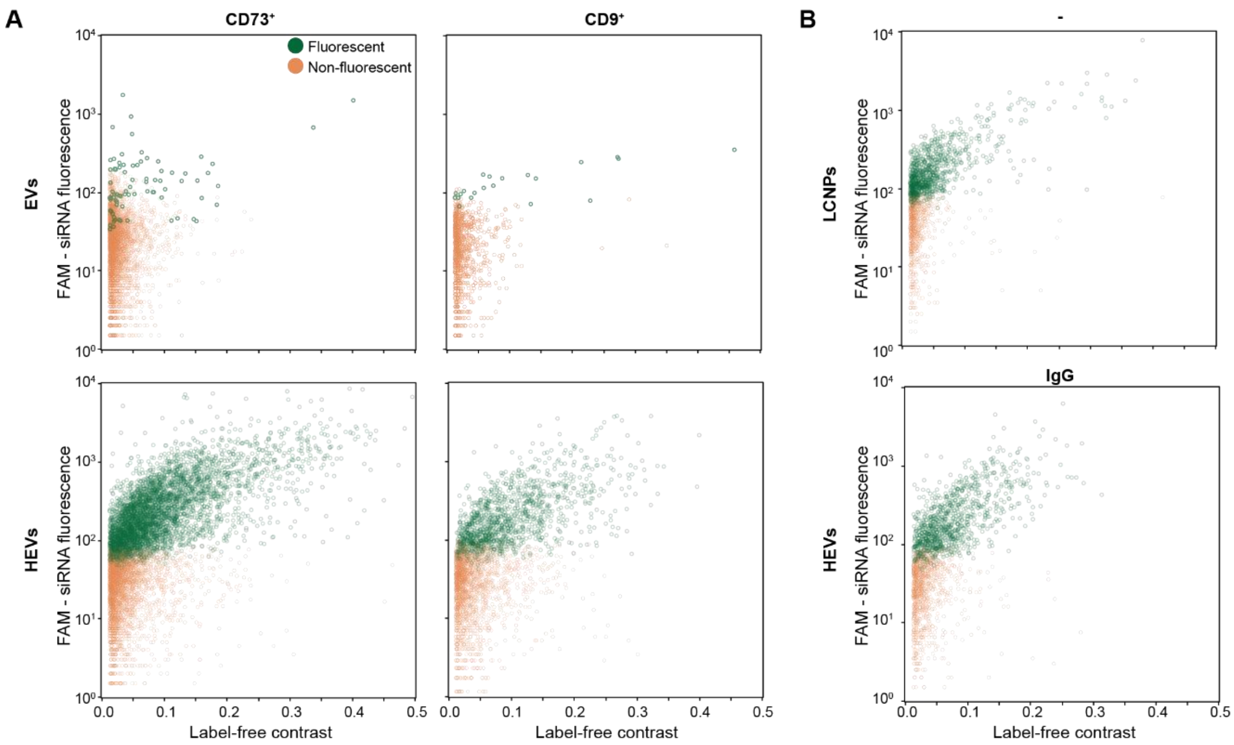


**fig. S11. Single-particle molecular fingerprinting and colocalization quantification of HEVs.** Native MSC-EVs mixed at a particle number ratio of 1:4 with FAM-siRNA-loaded (N/P 25) LCNPs after 30 min of incubation at 37 °C and pH7.4. (**A**) Representative bivariate dot plots for all the colocalized particle signals of EVs and HEVs immobilized during an immunoaffinity pulldown assay and expressing the respective target membrane protein (CD73 or CD9). The fraction of fluorescent (green) and non-fluorescent particle (orange) populations are indicated by their respective colors. (**B**) Representative bivariate dot plots of the fluorescent and non-fluorescent particle populations for LCNPs and HEVs bound non-specifically to the surface, respectively. For the sake of clarity, the number of particles in every bivariate plot has been down-sampled by a factor of 10.

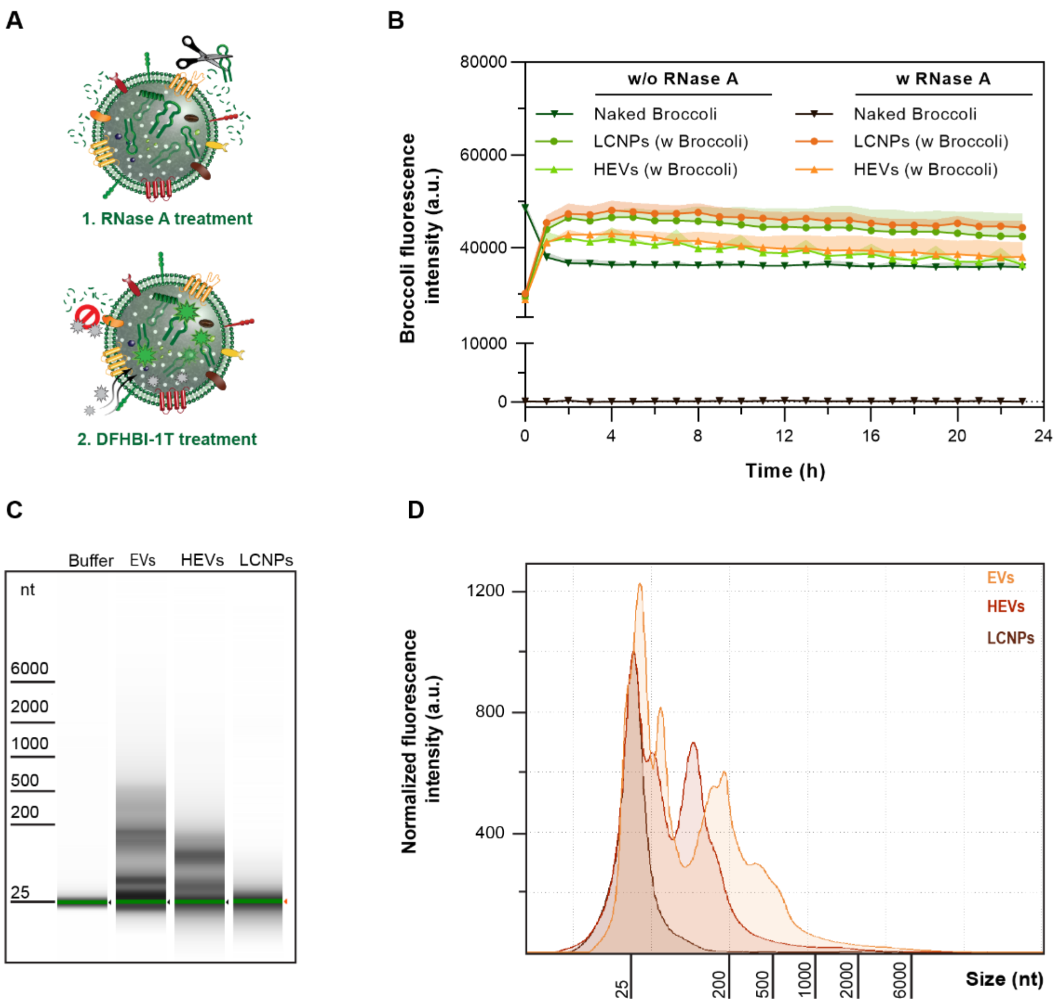


**fig. S12. Protection of exogenous and endogenous intraluminal NA cargo from nuclease degradation.**

(**A**) Scheme of the Broccoli RNA aptamer retention assay. Broccoli aptamer-loaded HEVs (fig. S9E) are treated with (1) RNase A (5 µg/mL) in the presence of the (2) aptamer-selective dye DFHBI-1T (20 µM). Broccoli that is released from HEVs/LCNPs will be degraded and is unable to interact with the dye. DFHBI-1T molecules can cross membranes and bind to intraluminal Broccoli protected from RNase-mediated degradation. The interaction of DFHBI-1T and Broccoli leads to an increase of the fluorescence signal *in situ*. (**B**) Broccoli fluorescence signal with or without RNase A assessed over 24 h at 37 °C. A total amount of 0.5 µg Broccoli loaded in HEVs/LCNPs and naked Broccoli are compared in the assay (n = 3). Data is expressed as mean + s.d. of n independent experiments. (**C**) Representative gel image from the TapeStation experiment performed with endogenous RNA isolated from EVs and HEVs that were hybridized with empty LCNPs at a particle number ratio 1:1 (pH 7.4 and 30 min at 37 °C). All formulations were incubated for 1 h at 37 °C with RNase A (5 µg/mL). Empty LCNPs alone are shown as the background control. (**D**) Overlay of the electropherograms from the TapeStation experiment.

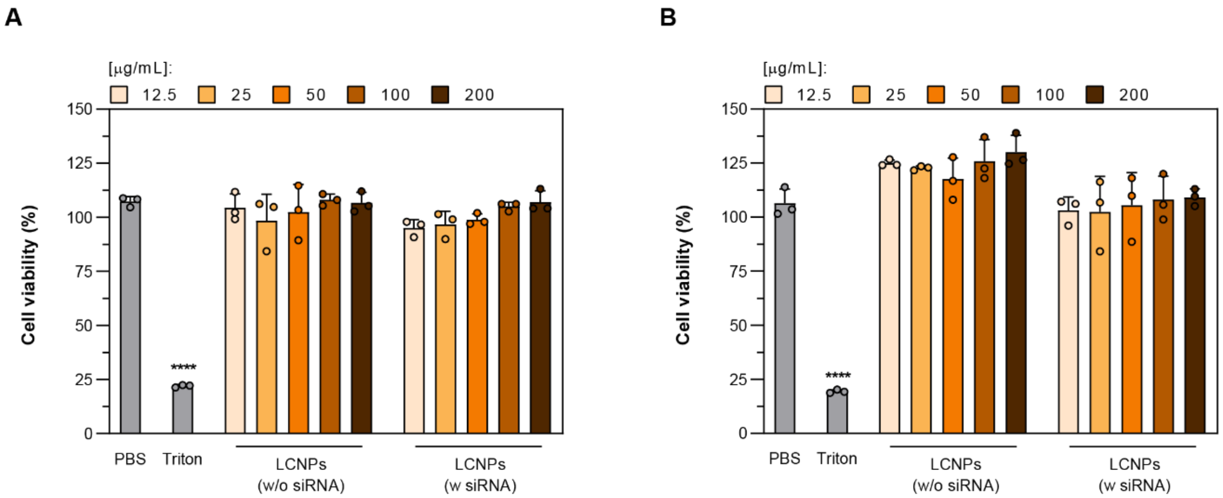


**fig. S13. HeLa cell viability after LCNP treatment.**

Cell viability was assessed after incubating various concentrations of LCNPs (from 12.5 to 200 µg/mL) with HeLa cells for (**A**) 4 and (**B**) 48 h (n = 3). LCNPs were either unloaded or loaded with siRNA (N/P 25). PBS and 1% (v/v) Triton™ X-100 were used as a negative and positive controls, respectively. Data is expressed as mean + s.d. of n independent experiments (n = 3). Statistical significance was calculated from an ordinary one-way ANOVA with Tukey's post-hoc test; ****$p < 0.0001$.

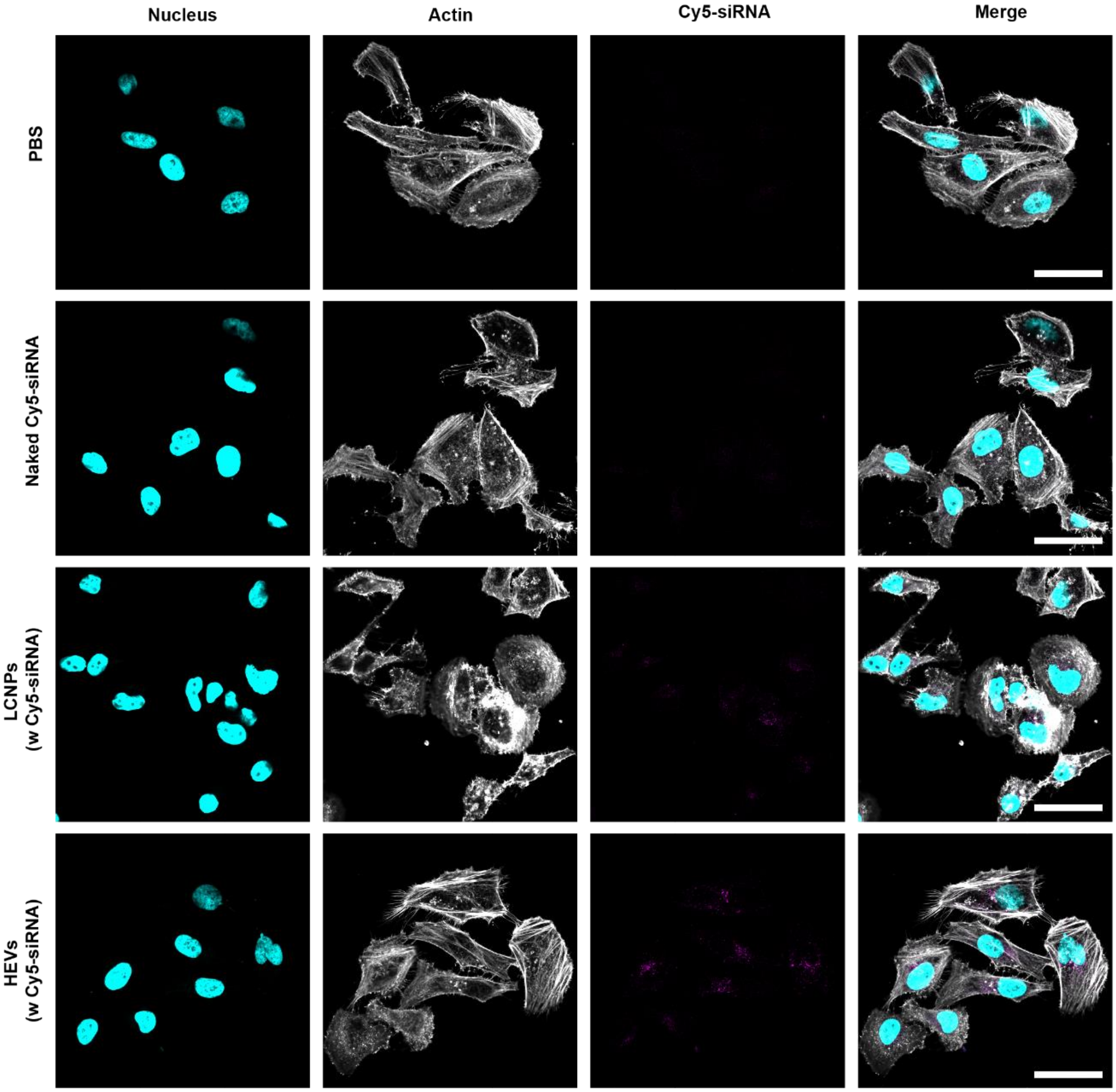


**fig. S14. Qualitative assessment of the uptake of Cy5-siRNA-loaded LCNPs and HEVs by confocal microscopy.**

The panel shows the three different imaging channels (from left to right): nucleus (cyan), actin (gray), Cy5-siRNA (magenta) and the merged image. HeLa cells were treated for 4 h with Cy5-siRNA-loaded LCNPs (N/P 25) and HEVs (100 nM Cy5-siRNA/well). Naked Cy5-siRNA and PBS serve as treatment controls. The confocal images corroborate the quantitative results from FACS analysis (Fig. 6A). Scale bars are 50 µm.

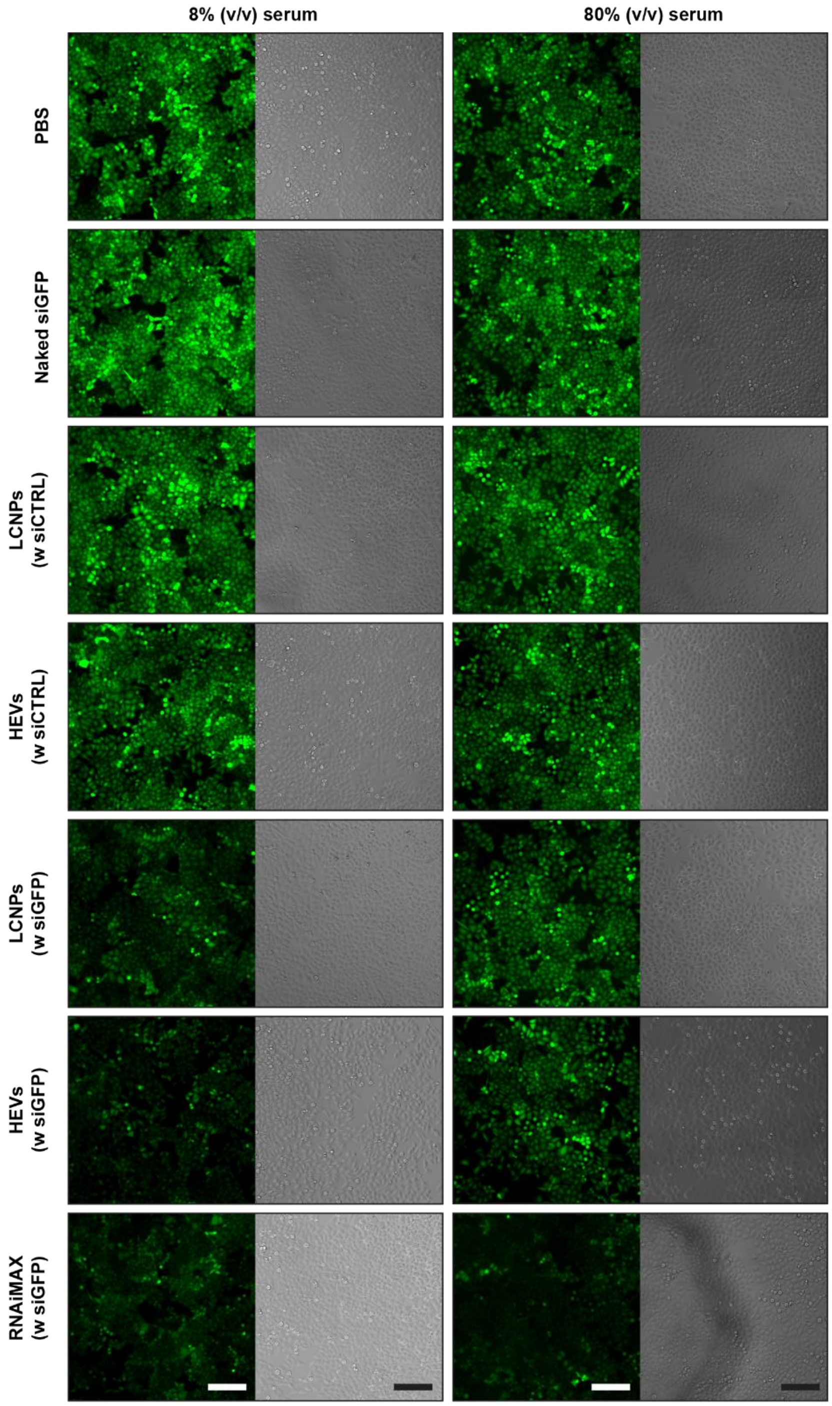


**fig. S15. Qualitative assessment of the GFP expression by live-cell fluorescence microscopy.** The panel pair shows the GFP fluorescence (left) and the phase contrast (right) images of HeLa-GFP cells treated with various siRNA-loaded LCNP (N/P 25) and HEV formulations (10 nM siRNA/well) and the RNAiMAX control at 8 and 80% (v/v) serum conditions. Images were taken 48 h after sample addition. The fluorescence images corroborate the quantitative results from FACS analysis (Fig. 7). Scale bars are 100 μm. The phase contrast image of the PBS-treated sample (80%, v/v serum) is slightly shifted.

**table S1. Characteristics of used NAs.**

| Identity | Length | Sequence | Molecular weight | Manufacturer |
|---|---|---|---|---|
| Broccoli RNA | 69 | GGGUUGCCAUGUGUAUGUGGGAGACGGUCGGGUC CAGAUAUUCGUAUCUGUCGAGUAGAGUGUGGGCUC | 22,680 (theor.) 22,721 (LC-MS) | Custom |
| GFP-siRNA | 21 | 5'-GAACUUCAGGGUCAGCUUGGG-3' (overhang sense: dGdG antisense: dTdT) | 13,365 | Custom by Microsynth AG |
| Control-siRNA | 21 | 5'-UAAGGCUAUGAAGAGAUACTT-3' (overhang sense: dTdT antisense: dTdT) | 13,270 | Custom by Microsynth AG |
| FAM-siRNA | 21 | ND | 13,854 | Sigma-Aldrich (# SIC007) |
| Cy5-siRNA | 21 | ND | 13,851 | Sigma-Aldrich (# SIC005) |
| FLuc-mRNA (5moU) | 1929 | ND | ~ 637,715 | TriLink BioTechnologies (# L-7202) |
| Cy5-FLuc-mRNA (5moU) | 2112 | ND | ~ 785,980 | Cellerna Bioscience (# 7001M) |

**table S2. List of antibodies for western blotting.**

| Name | Dilution (v/v) | Manufacturer |
|---|---|---|
| Goat anti-mouse IgG/HRP | 1:2000 | Dako Denmark A/S (# P0447) |
| Rabbit anti-goat IgG/HRP | 1:1000 | R&D Systems (# HAF017) |
| Mouse anti-human CD9 | 1:200 | Santa Cruz Biotechnology (# sc-13118) |
| Mouse anti-human CD63 | 1:500 | Santa Cruz Biotechnology (# sc-5275) |
| Mouse anti-human TSG101 | 1:200 | Santa Cruz Biotechnology (# sc-7964) |
| Mouse anti-human CD73 | 1:500 | Santa Cruz Biotechnology (# sc-32299) |
| Goat anti-human CD26/DPP4 | 1:200 | R&D Systems (# AF1180) |
| Mouse anti-human Grp94 | 1:200 | Santa Cruz Biotechnology (# sc-393402) |
| Mouse anti-human Calregulin | 1:500 | Santa Cruz Biotechnology (# sc-373863) |
| Mouse anti-human GAPDH | 1:1000 | Santa Cruz Biotechnology (# sc-47724) |

**table S3. Manual threshold settings for NanoFCM experiments.**

| Experiment | Threshold channels | | |
|---|---|---|---|
| | 488/10 | 525/20 | 670/30 |
| FAM-siRNA/CellTrace™ Far Red | 15 | 70 | 70 |
| Broccoli aptamer/CellTrace™ Far Red | 15 | 70 | 70 |
| Atto 488 DOPE/CellTrace™ Far Red | 10 | 25 | 115 |
| Cy5-mRNA/CFSE | 50 | 60 | 60 |
